# Properties of Two-Locus Genealogies and Linkage Disequilibrium in Temporally Structured Samples

**DOI:** 10.1101/2021.06.17.448867

**Authors:** Arjun Biddanda, Matthias Steinrücken, John Novembre

## Abstract

Archaeogenetics has been revolutionary, revealing insights into demographic history and recent positive selection in many organisms. However, most studies to date have ignored the non-random association of genetic variants at different loci (i.e., linkage disequilibrium, LD). This may be in part because basic properties of LD in samples from different times are still not well understood. Here, we derive several results for summary statistics of haplotypic variation under a model with time-stratified sampling: 1) The correlation between the number of pairwise differences observed between time-staggered samples (*π*_Δ*t*_) in models with and without strict population continuity; 2) The product of the LD coefficient, D, between ancient and modern samples, which is a measure of haplotypic similarity between modern and ancient samples; and 3) The expected switch rate in the Li and Stephens haplotype copying model. The latter has implications for genotype imputation and phasing in ancient samples with modern reference panels. Overall, these results provide a characterization of how haplotype patterns are affected by sample age, recombination rates, and population sizes. We expect these results will help guide the interpretation and analysis of haplotype data from ancient and modern samples.

## 1 Introduction

Multi-locus properties of genetic variation have been useful for studying evolutionary processes and maximizing the information extracted from population genetic data. Patterns of multi-locus variation are shaped by mutation *and* recombination events, generating novel combinations of alleles on chromosomes (i.e., haplotypes). The non-random association of alleles between two (or more) loci is known as linkage disequilibrium (LD) (Lewontin and Kojima, 1960; Hill and Robertson, 1968; Slatkin, 2008). Common measures of LD include the covariance and correlation in allelic state at two loci on the same haplotype within a sample (*D* and *r*^2^, respectively) (Hill and Robertson, 1968; Slatkin, 2008). The decay of LD as a function of the distance between genetic variants plays an important role in dating evolutionary events (e.g., Moorjani *et al.*, 2016), determining the accuracy of complex trait prediction (e.g., VilhjÁlmsson *et al.*, 2015), and moderating the power to map trait associated loci (e.g., Spencer *et al.*, 2009; Wray, 2005).

One approach for modeling variation at multiple loci has been through the use of coalescent theory (Kingman, 1982; Hudson, 1985). The coalescent process at multiple loci can involve both recombination (splitting events) and coalescence (joining events) of ancestral lineages, which means that there can be a different number of lineages at each locus at a given point in time (Hudson, 1985; Simonsen and Churchill, 1997; Durrett, 2008). Based on a two-locus coalescent model, Hudson (2001) developed a composite likelihood approach to estimate fine-scale recombination rates in early sequencing datasets. This initial approach paved the way for subsequent methods to estimate fine-scale recombination rates in humans, accommodating increasing model complexity (McVean *et al.*, 2004; Auton and McVean, 2007; Kamm *et al.*, 2016; Spence and Song, 2019). Also using a two-locus coalescent model, McVean (2002) was able to express metrics of LD in terms of properties of coalescent times. As the impact of changing demographic history on coalescent times is relatively straightforward, this advance enabled a more intuitive understanding of the impact of demographic history and sampling design on expected patterns of LD in data (McVean, 2002; Wakeley and Lessard, 2003).

A second major modeling framework for LD has been via “haplotype copying” models, such as the Li & Stephens’ model (Li and Stephens, 2003; Song, 2016). Haplotype-copying models provide a computationally efficient approximation for the likelihood of observed haplotype data generated with recombination (Fearnhead and Donnelly, 2001). As a result they have become a backbone of many analyses of population-genomic data, such as genotype imputation (e.g. Howie *et al.*, 2009), haplotype phasing (e.g. Loh *et al.*, 2016), and local ancestry inference (Price *et al.*, 2009; Lawson *et al.*, 2012).

In an increasing number of settings, samples are not all from the same time point. This is exemplified by the growing study of archaeogenetics, also known as ancient DNA (aDNA) studies (reviewed in Slatkin and Racimo, 2016; Llamas *et al.*, 2017; Skoglund and Mathieson, 2018). Archaeogenetic studies of humans have been able to reliably obtain genetic data from samples up to 45,000 years before present, although the majority of samples are from the past ~15,000 years (Skoglund and Mathieson, 2018).

For single locus data, genealogical models have been developed to quantify the impact of ancient samples on population genetic statistics, such as the expected site-frequency spectrum, the number of variants private to an ancient sample, and *F_ST_* (Rodrigo and Felsenstein, 1999; Forsberg *et al.*, 2005; Ortega-Del Vecchyo and Slatkin, 2018). In contrast, the impact of time-separation on patterns of linkage disequilibrium has not been fully explored.

Here we characterize patterns of haplotype variation in temporally stratified samples using a genealogical perspective. Analogous approaches for time-stratified samples in a coalescent framework have generally not been developed for the case of two or more recombining loci. One exception is the approach of Dialdestoro *et al.* (2016) that uses importance sampling over the space of latent ancestral recombination graphs when calculating the likelihood of observed sequence data for haplotypes at multiple time-points. Our work here contrasts to that of Dialdestoro *et al.* (2016) in that we obtain analytic solutions for two-locus scenarios and for the haplotype-copying model. The work presented here is complementary to previous work by Terhorst *et al.* (2015) who modeled how allele frequencies change for multiple loci using a Gaussian approximation to the Wright-Fisher model, though here we approach the problem from a coalescent perspective.

We focus on the case of two haplotypes at two loci because it represents the simplest case of time-stratified sampling across multiple loci, is analytically tractable compared to larger sample sizes, and provides insight on expected patterns in data (Hudson, 1985; McVean, 2002). We first show how time-stratified sampling affects the joint properties of genealogies at two loci, demonstrating that the time gap between a pair of samples has an impact on the rate of decay in the correlation of genealogical statistics and corresponding patterns of variation with recombination distance. We also analyze the behavior of fitting the haplotype copying model with samples of different ages, in particular when the test haplotype is from a time-point in the past compared to a modern haplotype panel. Overall, our results show the effect of time-stratified sampling on expected patterns of haplotypic variation, and their implications for the further development of population genetic methods.

## 2 Results

### 2.1 Two-Locus Genealogical Properties

To model two haplotypes at two loci with time-stratified sampling, we adapted a previously developed continuous time Markov process for modeling ancestral lineages at two loci (Hudson, 1983, 1990; Simonsen and Churchill, 1997). The states in the model are triplets (e.g., (2, 0, 0)) that depict the number of lineages ancestral to both loci, locus 1, or locus 2, respectively. Coalescence and recombination events eventually lead to an absorbing state where *both* haplotypes have coalesced at *both* loci (the state (1, 0, 0), Figure A1). Analytical results for joint moments in the coalescent times in this model have been previously obtained for the case where samples are taken at the present (Hudson, 1983; Simonsen and Churchill, 1997; Durrett, 2008, Chapter 3).

Here, to analyze the case of time-stratified sampling, we assume that one of the haplotypes has been sampled at time *t_a_* in the past (in coalescent units) and the other at the present. With this time gap in sampling, there are two natural phases in the ancestral process: (1) the time between the present and when the ancient haplotype is sampled (*t < t_a_*), and thus only the lineage of the modern haplotype can evolve at each locus, and (2) the time when the lineages of both haplotypes (modern and ancient) are evolving through the full state space of the ancestral process (*t* ≥ *t_a_*).

For this two-phase ancestral process, we derived expressions for the covariance between the *T_MRCA_*’s at two loci (*A* and *B*), as well as the total branch lengths (*L_A_*,*L_B_*) separated by a population-scaled recombination distance, *ρ* = 4*N_e_r*, where *r* is the per-generation probability of recombination.

The derivation proceeds by recognizing that a key aspect of the two-phase process is the effect of recombination during the first phase, when only the modern lineage is evolving backwards in time (*t < t_a_*, see Appendix A1). During this phase the process has only two states, “uncoupled” and “coupled”. By “uncoupled” we mean that the ancestral lineages are evolving independently at each locus, whereas “coupled” means that they are evolving as a joint ancestral lineage. The starting state for the second phase of the ancestral process (when *t ≥ t_a_*) is either that the modern haplotype’s ancestral lineages are coupled at both loci or uncoupled from one another. We obtain the time-dependent probability of being in the uncoupled state by exponentiating the 2 × 2 rate matrix **Q** for the reduced state-space of the ancestral process during *t < t_a_*, 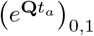, where 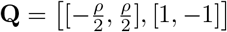. By doing so and taking different limits, we find:

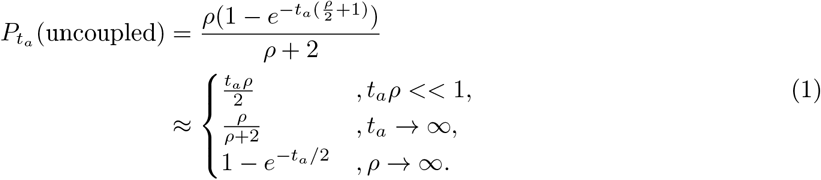

Figure 1B shows for either large time-separation (*t_a_*) or large population-scaled-recombination rates (*ρ*), it becomes more likely that the modern haplotype is in the uncoupled state by the time the process encounters the ancient haplotype. Since the remaining dynamics are the same as the two-locus ancestral process with two contemporaneously sampled haplotypes, we thereafter leverage known results for the two-locus ancestral process (Simonsen and Churchill, 1997; McVean, 2002; Durrett, 2008, Chapter 3). In the next two sections, we take this modeling approach to derive the expectations of observable quantities from time-staggered haplotype data.

**Figure 1:**
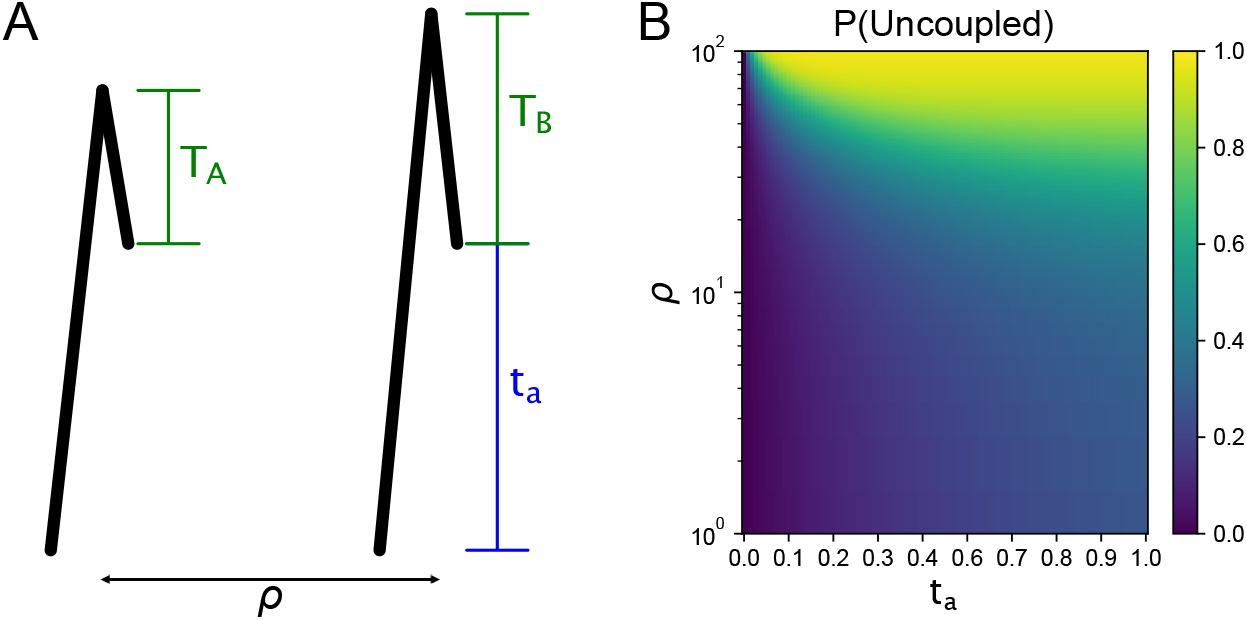
**(A)** Schematic of genealogies at two loci separated by a population-scaled recombination distance *ρ* (*ρ* = 4*N_e_r*). The parameter *t_a_* represents the sampling time of the haplotype (measured in coalescent units, i.e., scaled by 2*N_e_*). The random variables *T_A_* and *T_B_* are the additional time to coalescence at locus *A* & *B*, after *t_a_*. **(B)** The probability of the modern haplotype being “uncoupled” at the time of ancient sampling as a function of *t_a_* and *ρ*. In this setting, “uncoupled” means that the ancestral lineages at locus *A* and *B* are not on the same haplotype, enhancing the probability of different *T_A_* and *T_B_* occurring at each locus.

### 2.2 Correlation in pairwise differences

The number of pairwise differences between two haplotypes at each of two loci is an observable summary of genetic variation at linked loci in time-sampled sequence data. To investigate the properties of the joint distribution on pairwise differences at two loci (locus *A* and *B*), we continue to assume a model with recombination occurring at a rate *ρ* between them and no recombination occurring within each. For each locus, as is typical in coalescent models, we assume an infinite-sites model with mutations arising on each lineage as a Poisson process with rate 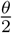, where *θ* = 4*N_e_μL*, *μ* is the per-basepair per-generation mutation rate, *L* is the size of the locus (in basepairs), and *N_e_* is the effective population size.

Following the approach described in the preceding section, we derive the correlation of pairwise differences for the case with time-stratified sampling (see Appendix A1). In particular, we use the fact that the correlation in the number of pairwise differences at locus *A* and *B* can be expressed in terms of the correlation in the total branch length between the loci (Wakeley and Lessard, 2003; Hobolth *et al.*, 2019). We find the correlation in pairwise differences between two loci to be:

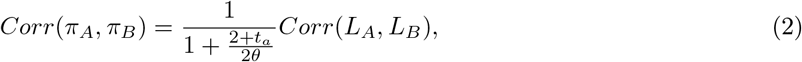

where *Corr*(*L_A_, L_B_*) is the correlation in total branch length at locus *A* and locus *B*. In Appendix A1 (building on previous results from Hudson, 1983; Simonsen and Churchill, 1997; Durrett, 2008, Chapter 3), we derive its exact form and several limiting values to be:

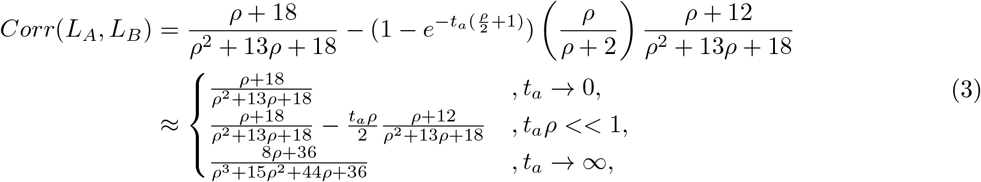

As the equations show, the correlation in pairwise differences is affected by the age of the ancient sample *t_a_* in two ways. The first effect is due to the factor in Equation 2 that decreases as *t_a_* increases and is not dependent on *ρ*. The second effect occurs in how *t_a_* affects *Corr*(*L_A_, L_B_*) (Figure 2A). For values of *t_a_ρ <<* 1, the correlation decays linearly with *t_a_* and with 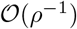 for *ρ*. The decay decreases more rapidly as 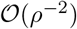 when *t_a_ρ >>* 1 and as *t_a_* gets large (the third case in Equation 3). This is because of the additional time (*t_a_*) that the recombination process has to break apart the shared genealogical history at each locus.

**Figure 2:**
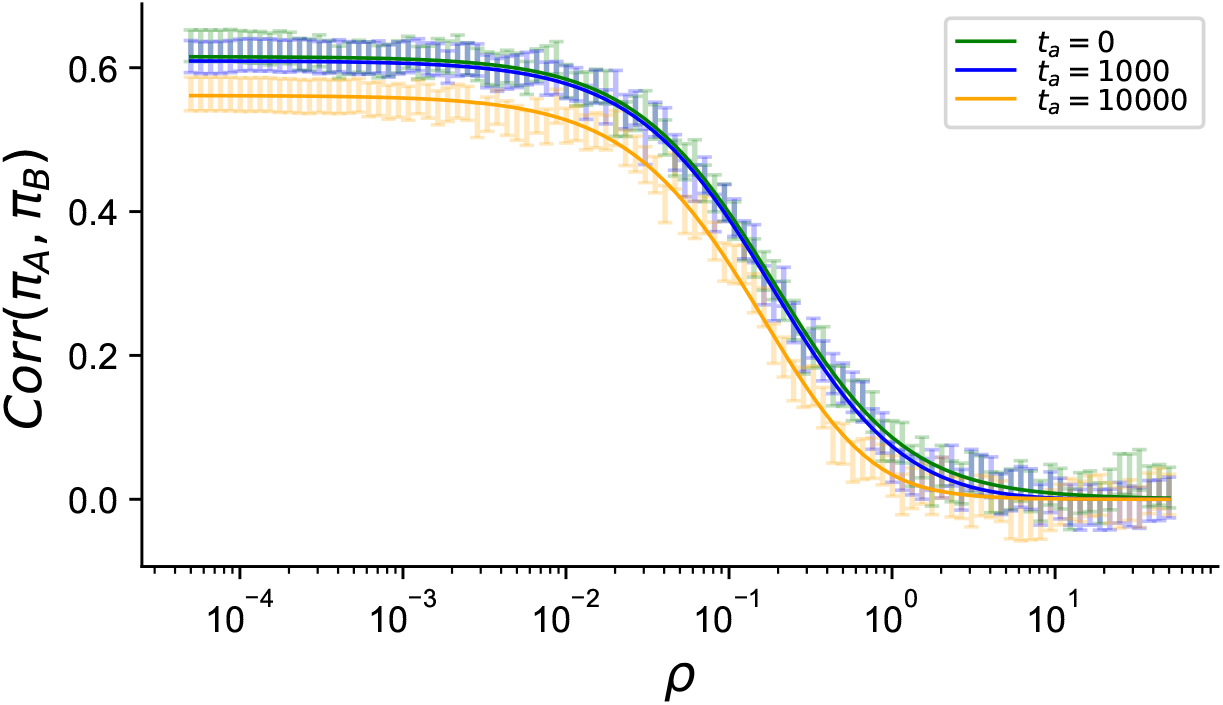
Theoretical (solid lines) and simulated correlation between pairwise differences in a constant-size demography (*N_e_* = 10^4^) at different sample ages (in generations). Comparison of theoretical prediction of *Corr*(*π_A_, π_B_*) with data from two-locus coalescent simulations with *θ* = 0.4 (see Methods). Solid blue and orange lines are the theoretical predictions for *Corr*(*π_A_, π_B_*) from Equation 2

#### 2.2.1 The impact of non-equilibrium demographic history on the correlation in pairwise differences

To explore the effects of varying population size through time, we simulated haplotype data under models of constant size, instantaneous growth, and trajectories inferred from previous studies of human populations that include both bottlenecks and growth (Tennessen *et al.*, 2012; Browning and Browning, 2015) (Figure 3). Motivated by how most human ancient DNA data are from approximately the last 15,000 years, we investigated the correlations on a timescale of 500 generations.

**Figure 3:**
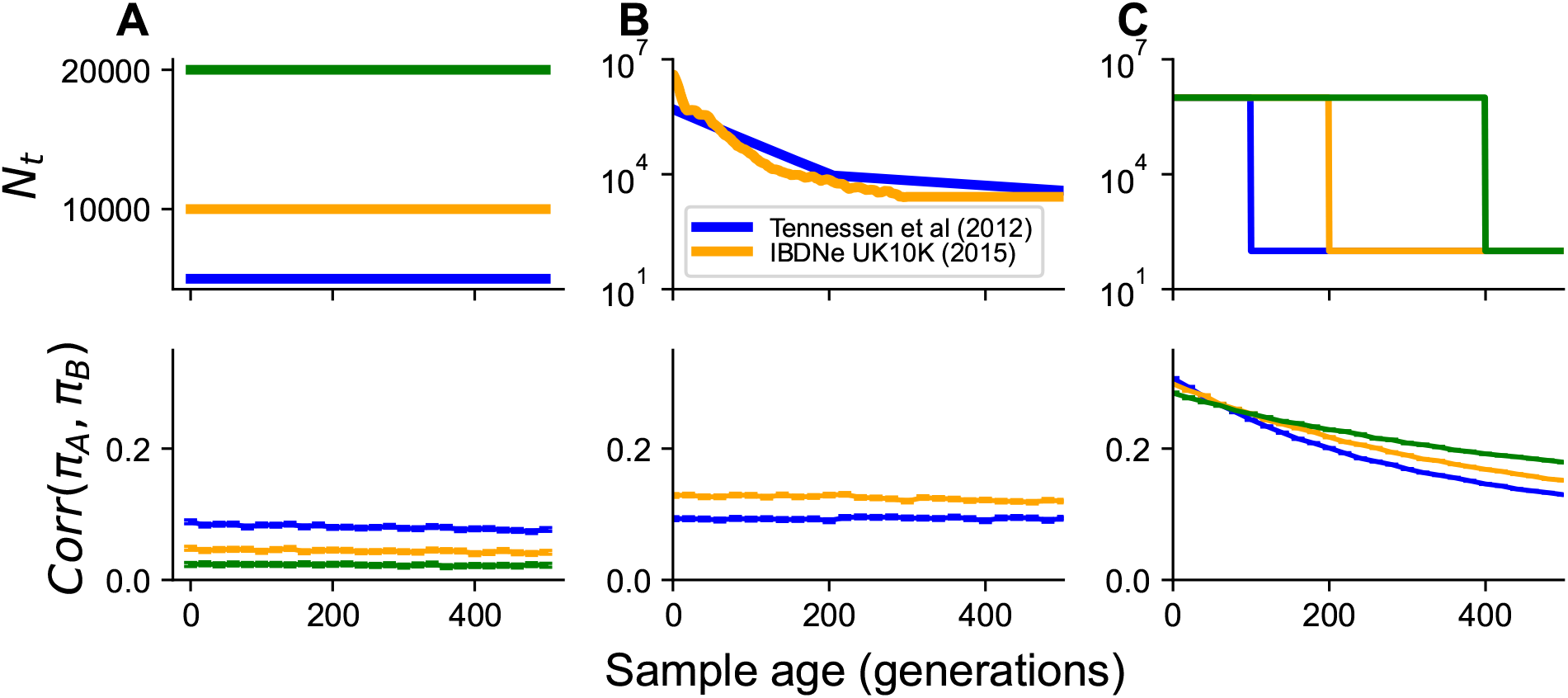
The impact of varying demographic history on the correlation in pairwise differences at two loci. For all simulations, the recombination rate between the loci was set to 10^−4^ per generation (approximatley 10kb, assuming 1 cM per 1Mb). Simulated scenarios include: **(A)** constant population size, **(B)** inferred models of population growth, and **(C)** models of instantaneous population growth. Each timepoint had 50000 replicate simulations.

In models with constant population size, larger population sizes lead to smaller inter-locus correlations (lower LD). In all our simulations *ρt_a_* << 1, so on the time-scale of 500 generations, the correlation in branch length decreases linearly as expected with sampling age (Equation 2, Figure S2A). Across all population sizes we observe significantly negative relationships between sample age (on the coalescent scale) and the correlation in branch length akin to what we predict in Equation 2 (for linear regression of *Corr*(*L_A_, L_B_*) ∼ *βt_a_*, we find for 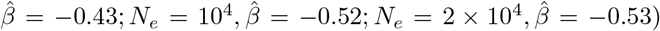. The negative effect of *t_a_* on the correlation in total branch length in turn decreases the correlation in pairwise differences (Figure 3A).

When simulating under the population size trajectories from Tennessen *et al.* (2012) or from the Browning and Browning (2015, “UK10K IBDNe model” in reference to the original dataset), the correlations are larger for the UK10K IBDNe model, which includes a larger population size in the last few generations but an overall *N_e_* (estimated using Watterson’s estimator, see Methods) that is smaller than the Tennessen model (*N_T ennessen_* ≈ 6922.91; *N_UK_*_10*K*−*IBDNe*_ ≈ 2670.19) (Figure 3B). In a linear model, the correlation in pairwise differences decreases with age under the UK10K IBDNe model (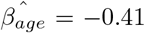, 95% CI=[0.51, 0.31]) and not in the Tennessen *et al.* (2012) model (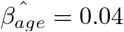, 95% CI: [−0.03, 0.12]).

For the case of step-wise population growth, (Figure 3C) we make three observations. First, the decrease in the correlation in pairwise differences is no longer approximately linear with time but decays non-linearly, with the rate of decay decreasing with sample age. Second, the correlation in pairwise differences is highest at short time-scales for the most recent growth event, and at long-timescales for the most ancient growth event. This can be interpreted again as a result of the very low *N_e_* in this setting such that the factor scaling the correlation in pairwise differences (Equation 2) dominates the behavior after *t_a_* ≈ 150 generations (when the correlation in branch length is similar across all settings). Third, the correlation in the branch length is substantially higher (*>* 0.8) when compared to the previously inferred demographies (Figure S2)

The step-wise growth scenario is interesting in that due to the large, recent increase in population size, we expect roughly star-like genealogies with coalescent times concentrated around the start of the growth event (Slatkin, 1996; Rosenberg and Hirsh, 2003). In this scenario, we find the correlation between loci in the branch lengths is increased greatly (Figure S2C, Figure S7) which contributes to elevating the *Corr*(*π_A_, π_B_*). At the same time, as *θ* is decreased relative to other scenarios (due to lower *N_e_*), we do not see as drastic an increase in the correlation between pairwise differences as in the branch length (Equation 2).

Finally, we also investigated the correlation in pairwise differences in a two-population model of divergence without gene flow. We assume the modern and ancient haplotype are each sampled from different populations. In this scenario, *both* the ancient and modern haplotypes can become uncoupled prior to any possibility of inter-haplotype coalescence lowering the expected correlation in pairwise diversity (Appendix A1.1). In this model, we find the correlation in number of pairwise differences decreases as a function of the sum of the divergence time and the sampling time (*t_div_* + *t_a_*) (Figure S1).

#### 2.2.2 Correlation of pairwise differences in time-staggered whole-genome sequencing data

Next, we explored the correlation of pairwise differences in modern and ancient human whole-genome sequencing data with two high-coverage samples from two different ages. We restricted to analyzing high-quality whole genome sequencing data to avoid ascertainment biases and to more accurately estimate pairwise differences (see Methods, Figure 4).

**Figure 4:**
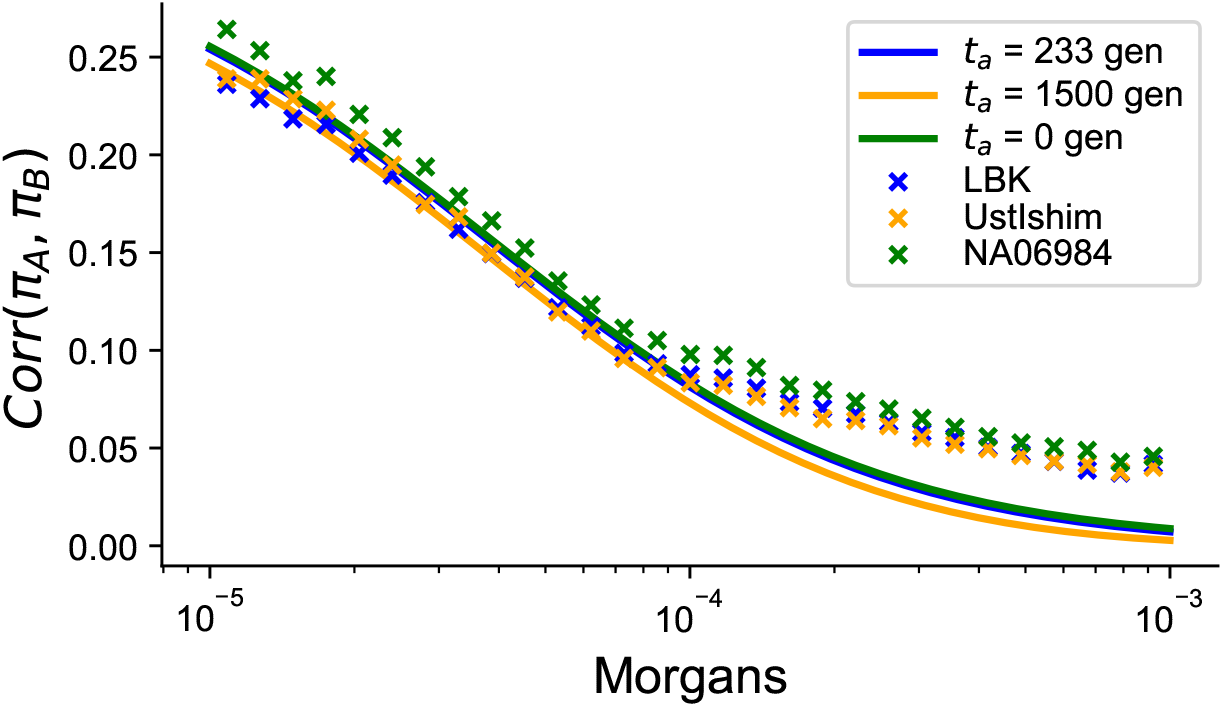
Comparison of the correlation in pairwise differences between LBK, Ust-Ishim, and a modern CEU control individual. Points represent the Monte-Carlo estimate of the pairwise correlation between randomly chosen pairs of loci (see Methods). When computing the theoretical curves, we used *N_e_* = 10^4^ and a mutation rate *μ* = 1.2 × 10^−8^ per basepair-per-generation.

The first sample we chose is an approximately 7000 year old sample from modern-day Germany associated with the Linear Ban Keramic culture and labeled variously in previous studies as the Stuttgart *LBK* sample or simply the *LBK* sample (Lazaridis *et al.*, 2014). The second sample is approximately 45000 years-old and from Western Siberia, labeled *Ust-Ishim* (Fu *et al.*, 2014). These samples to have an order of magnitude difference in the sampling time-scale (thousands vs. tens-of-thousands years).

To complement the theoretical analysis, we calculate the correlation of *π_A_* and *π_B_* in two samples across a large number of paired 1 kilobase windows (see Methods). We find that the correlation in pairwise differences as a function of recombination distance for Ust-Ishim is not significantly different from LBK when only relying on a single control individual (Fig. 4, Binomial Test, fraction of pairs of loci *Corr*(*π_A_, π_B_*)_*LBK*_ > 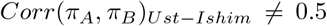, *p* = 0.58466). When repeating the hypothesis test using a modern CEU individual (’NA06984’) and comparing against the Ust-Ishim sample we found that using a modern control individual we were able to reliably reject the null hypothesis of no difference (*p* = 0.016; Binomial Test). This result also holds across numerous different choices of the modern control individual and when taking the mean estimate of the correlation across multiple control individuals as shown in Figure 4 (*p* = 0.361 vs. *p* = 1.86 × 10^−9^; Binomial test), suggesting that the correlation in pairwise differences has greater power to differentiate between modern and ancient samples rather than comparing two ancient samples of different ages.

Qualitatively, we find that there is also a lack of fit with the theory derived for constant population size, in that we observe at longer recombination length scales an excess of correlation in segregating sites. We speculate that this may be due to the effects of population bottlenecks or recent growth in the history of non-African populations leading to elevated levels of linkage disequilibrium at this scale that is not well-captured by our theoretical model of constant population size (Reich *et al.*, 2001; Kamm *et al.*, 2016; Ragsdale and Gravel, 2019) (Figure S3).

### 2.3 Linkage disequilibrium with time-stratified sampling

To directly relate the joint genealogical properties described above to patterns of LD, we investigated the normalized expected product of linkage disequilibrium (*D*) between the ancient and modern samples:

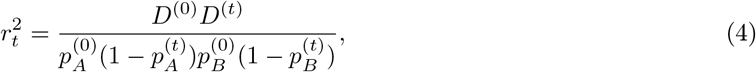

where 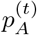 is the frequency of the derived allele at the first locus at time *t* and 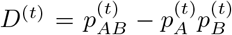 is a classic measure of LD in the sample of individuals from time *t* (Lewontin and Kojima, 1960).

Using the genealogical identity coefficients from McVean (2002), we derive the ratio of the expectations of the product of linkage disequilibrium between time-points. Motivated by arguments put forth by Ragsdale and Gravel (2019) and McVean (2002) that express statistics of LD by taking the ratio of expectations (i.e. 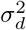), we similarly take the ratio of expectations of 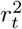 in Equation 4 to derive a time-stratified analog of 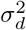. In Appendix A3, we derive an expression for the joint product of LD across both timepoints 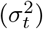:

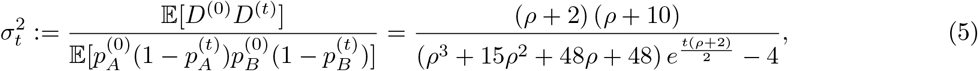

When *t* = 0, Equation 5 reduces to the expression for 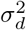, as shown in McVean (2002). Both simulations and our theoretical predictions show that larger time-separation between samples qualitatively decreases the joint product of LD (Figure 5).

**Figure 5:**
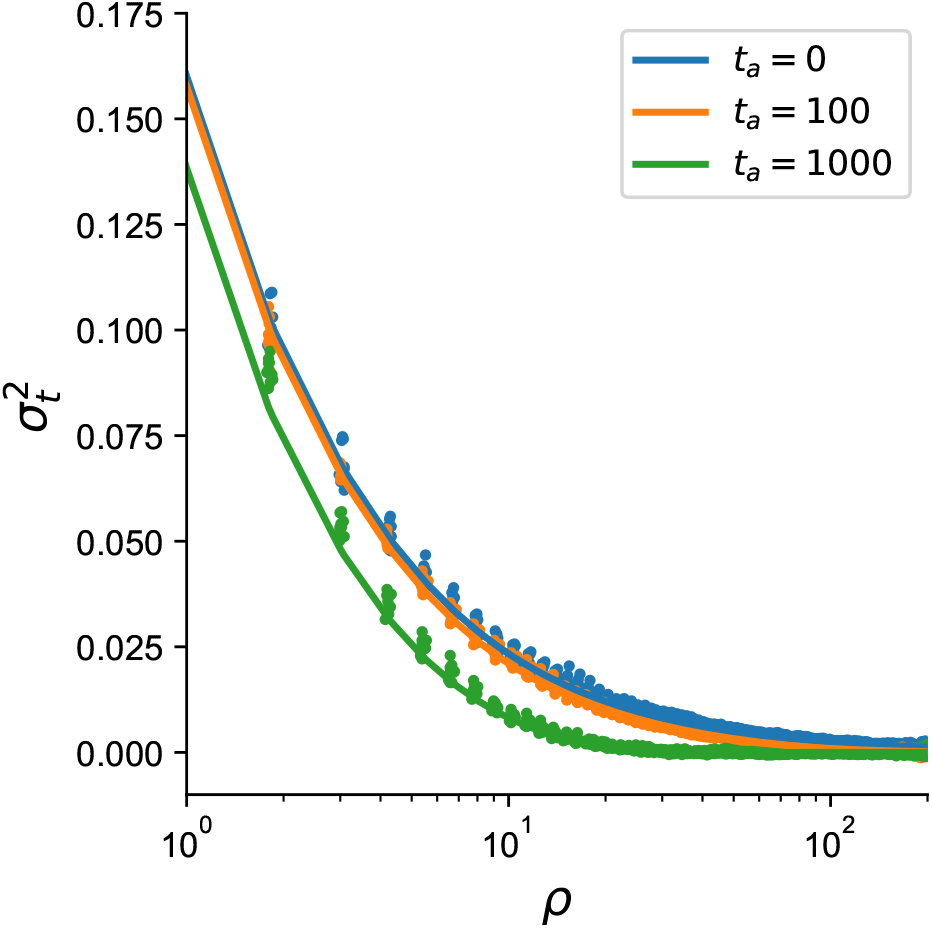
Joint product of linkage disequilibrium between samples separated by *t_a_* generations across different population-scaled recombination rates *ρ* (see Methods). Dots represent results from simulation and solid lines are theoretical predictions from Equation 5.

### 2.4 The impact of time-stratified sampling in haplotype-copying models

We next consider the scenario where one would be interested in modeling an ancient haplotype as a mosaic of modern haplotypes, as might arise when trying to phase or impute ancient DNA genotypes using a reference panel of modern haplotypes and the popular Li & Stephens haplotype-copying model (Li and Stephens, 2003; Song, 2016). We specifically use a modified model where the recombination map positions are known *a priori* (see Methods) (Lawson *et al.*, 2012). We focus on the maximum-likelihood estimate of the haplotype copying jump rate (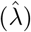) for a given test haplotype as it copies off the reference panel. We view 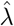 partly as a summary statistic reflecting the length scale of copying tracts and as an indicator of the expected accuracy of imputation (Stephens and Scheet, 2005; Jewett *et al.*, 2012).

The time-separation between the ancient haplotype and modern sample provides an opportunity for recombination events to occur among the modern reference haplotypes before the ancient lineage is able to coalesce with any individuals from the modern panel (Equation 1, Figure 1). Thus, we expect higher jump rates as the sample age *t_a_* increases. We also expect coalescence *within* the modern panel will contribute to higher jump rates with increasing *t_a_* by effectively reducing the panel size moving farther back in time.

Using the first time to coalescence between the ancient target and a member of the modern panel, we observe a saturation effect when increasing the modern panel size (Appendix A4, Figure S4). The time until the first coalescent event involving the ancient sample is equal to the length of the external branch in the local genealogy that leads to the ancient sample, and affects the rate of recombination events that can induce switch events in the copying model. The time to the first coalescent involving the ancient sample and the modern panel decreases as a function of the reference panel size, *K*. However, as the age of the sample increases, the number of lineages extant to the reference sample become smaller, making the time to first coalescent event more similar across modern reference panel sizes.

Using simulations with populations of constant size, we find that the realized copying jump rate indeed increases with age, and does so monotonically as a function of the age of the test haplotype under a model of constant population size (Figure 6A). The simple monotonic relationship can break down in non-equilibrium demographic models. For instance, in demographic models including recent population growth we find that there is an initial decrease in 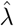 from the present to approximately 150 generations ago before a more rapid increase moving back into the past (Figure 6B) (Browning and Browning, 2015; Tennessen *et al.*, 2012). A similar results is observed more dramatically in simulations of instantaneous growth, with a common feature being a decreasing relationship between 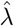 and sample age up to the time of onset of the instantaneous growth (Figure 6C).

**Figure 6:**
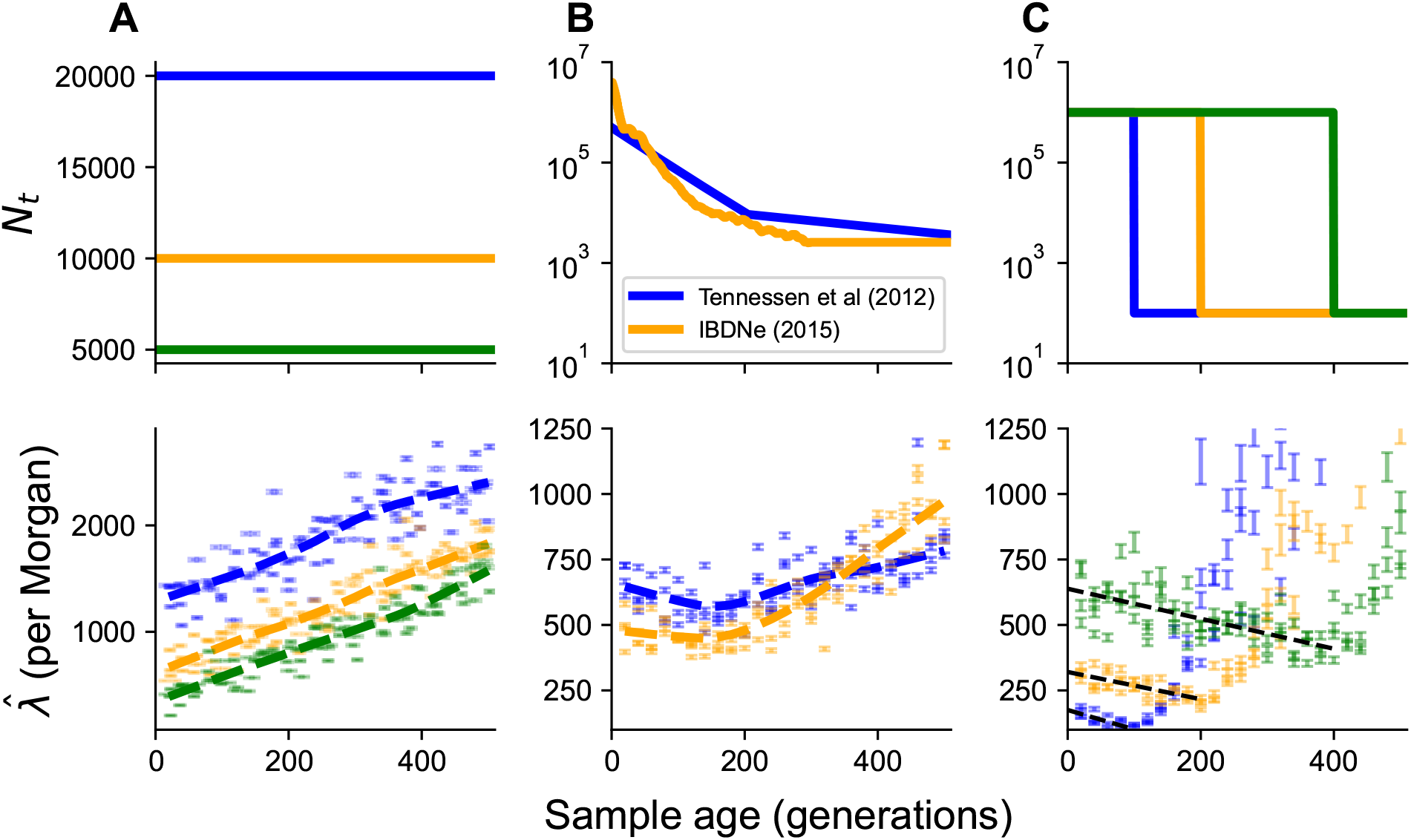
Estimation of haplotype copying jump-rate against sample age for different models of population demographic history (top row). **(A)** Constant population size, **(B)** previously inferred models of recent population growth, and **(C)** models of instantaneous population growth. The inferred parameters should be interpreted in terms of the average jumps per Morgan.

#### Haplotype-copying jump-rates in human ancient DNA data

To compare our simulation experiments on the dependence of the jump-rate with sampling time to empirical data, we applied our jump rate estimation to a collection of 1159 ancient human samples (see Methods). To avoid potential errors introduced by statistical phasing, we analyzed only haploid carriers of the X chromosome by taking samples labeled as male in both the ancient data and the modern reference panel (Thousand Genomes Project data, Auton *et al.*, 2015). Thus, the analysis used 47, 094 bi-allelic SNPs observed on the X chromosome. To avoid the potential effects of population structure confounding the impact of time-stratified sampling and to maximize the sample size, we focus primarily on Europe as it is the region with the highest density of aDNA samples, and we used *n* = 49 CEU male X chromosomes to define the modern reference panel (see Figure S5 for experiments with alternate panels).

Based on copying jump rates estimated across 344 ancient male X-chromosome samples from across Europe (see Methods for a description of the dataset), we find that the estimated jump rate *decreases* as a function of sample age (Figure 7A). Accounting for spatial variables (Latitude, Longitude, and Latitude × Longitude) in a linear model (see Methods, Figure S5), we find the effect of sample age on the estimated copying jump rate is negative and statistically significant (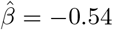; 95% CI=[0.63, 0.46]). Filtering for the highest 25% coverage individuals did not change the result (Figure S8). The inferred haplotype copying error rate (*E*) also decreases with age, suggesting the observed decrease in *λ* is not an artifact of the inference procedure (Figure S9).

**Figure 7:**
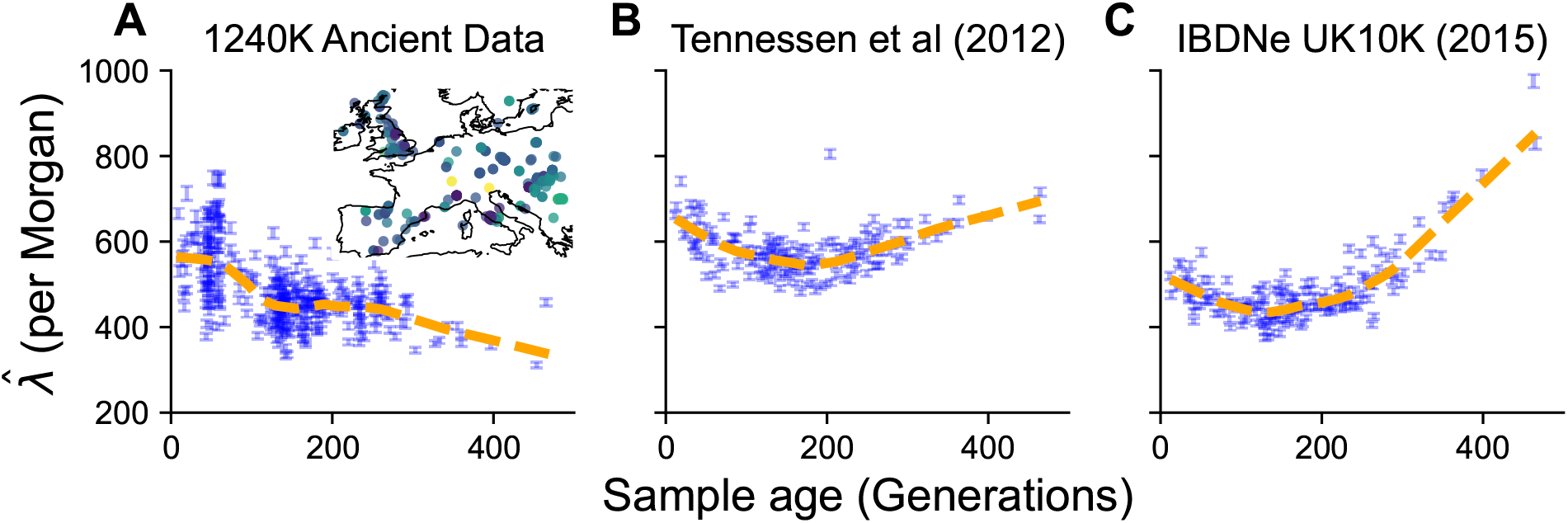
Comparison of estimated haplotype copying jump rates between real data and simulations. **(A)** Estimate of the jump rate in ancient male X-chromosomes within 1500 kilometers of central Europe. **(B)** Maximum likelihood estimates of the haplotype copying jump rate using simulated X-chromosomes under the model of Tennessen *et al.* (2012).**(C)** Estimated jump rates using simulated data under the model of Browning and Browning (2015).

This decrease is contrary to our idealized simulations with constant population size (Figure 6) and in agreement with the simulations involving some aspect of recent growth (Figure 6B,C). To make the comparison more exact, we replicate simulations of Tennessen *et al.* (2012) and Browning and Browning (2015) with the exact temporal sampling structure of the real 344 samples and using a sex-averaged recombination map for the X chromosome (Kong *et al.*, 2010). With these simulations, we are able to replicate an initial decrease in the jump-rate as a function of sampling time (Figure 7B,C). However, the simulations do not capture the duration of the decrease in jump-rate with sample age, which we find to be ≈ 400 generations in the real data.

## 3 Discussion

In this article, we have developed theory to understand the effects of serial sampling on patterns of haplotype variation in the context of two models, the two-locus coalescent model and the haplotype copying model. We primarily focused on these models because they provide theoretical results for the expected patterns of linked variation and they underlie standard approaches to analyze modern haplotype data.

We find that with larger time-separation between samples, the correlation in branch length at two loci decreases by an amount proportional to the probability of uncoupling of a sampled modern haplotype over *t_a_* units of time (Equation 2). In constant-size populations and small values of *t_a_ρ*, the decrease is linear in time. As *t_a_* increases the decay of correlation happens as 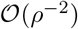 vs. 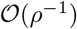. Intuitively, the additional marginal branch length on which a recombination event can occur (2 + *t_a_* vs. 2 in expectation) is disrupting between locus correlation. For larger values of *t_a_* there is an additional decrease in correlation that arises from the impact of mutations (the denominator of Equation 2). For small values of *t_a_* (*t_a_ <<* 2 coalescent units) the correlation of branch length essentially determines the behavior of the correlation in observable number of differences between two loci. The analysis of the expected behavior of the joint LD coefficient between data sampled at different times also showed that ancient sampling decreases the associations between loci, leading to a systematic decrease in the joint product of LD across all recombination scales (Figure 5).

Our analysis of the haplotype copying rate 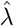 revealed interesting impacts of demographic history. In constant-size models, the inferred copying rate increased with the sample age as one might expect due to recombination events; jowever, in cases of strong recent population growth (Reppell *et al.*, 2014; Tennessen *et al.*, 2012; Browning and Browning, 2015) the inferred copying rate decreases initially with age and then increases. To understand this, consider how the haplotype-copying jump-rate, 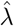, is inversely related to the expected branch-length shared between an ancient haplotype and a member of the modern panel, because recombination events that occur on these branches can initiate copying-switch events (Paul *et al.*, 2011; SteinrÜcken *et al.*, 2013; Li and Stephens, 2003). In cases with rapid population growth, there are initially limited numbers of coalescent events, followed by a high rate when the population is small, looking backwards in time. Samples that are sampled sequentially closer to the onset of growth have shorter branch length on which potential switch events occur, producing the initial negative relationship. For samples that are sampled *more ancestrally than* the onset of population growth, we find that the jump rate increases as the coalescent time are no longer affected by the onset of growth (Figure 6C).

The possibility of a negative relationship between copying rate and sample age was supported in our empirical analysis of ancient DNA data from western Eurasia. However, the empirical data from western Eurasia showed a more extended decrease in the jump rate, reaching over ~400 generations, relative to what was seen in the demographic models simulated. There are two potential types of explanation for the discrepancy between the simulations and observations: an artifact of the aDNA data or an aspect of demography that is not captured well by the existing models used for simulation. Regarding the aDNA data, in our empirical analysis, we do not find any significant effects of coverage on the qualitative result that the jump-rate decreases as a function of time (Figure S8B). If error rates increase with sample age it would seem to run counter to the observed result, causing elevated jump rate estimates as one goes further back in time, however this is not what we observe in our joint estimation (Figure S9). Some complex form of reference bias increasing with age and interacting with the haplotype copying model may be plausible. Regarding explanations based on the demographic models, the duration of the decrease in estimated copying rate could potentially be due to smaller local population sizes in the more distant past than is reflected in the models. This is particularly relevant given the time-scale of ~400 generations (~12, 000 years) as this extends into the Mesolithic and Paleolithic eras during which populations were likely small in overall size and deeply structured (Premo and Hublin, 2009; Haak *et al.*, 2015; Skoglund and Mathieson, 2018). If ancestral population structure existed in this period it may have biased inferred effective population size upwards in models that were fit under the assumption of a single panmictic population (Li and Durbin, 2011, Section S1.6). Overall, the result suggests there may be interesting insights to be gained by more detailed empirical analyses of haplotypic patterns in ancient DNA.

Many methods have been developed in the context of haplotype copying models, from imputation and phasing (Howie *et al.*, 2009, e.g.,), estimation of recombination rates (Li and Stephens, 2003, e.g.,), to fine-scale ancestry estimation (e.g., Lawson *et al.*, 2012). Our theoretical results leave important considerations for each of these application domains with serially-sampled data. For imputation and phasing, the increase in the copying jump rate as a function of time under constant population sizes implies that linkage disequilibrium will be lower in relation to the first coalescent time with a member of the modern panel, and will lower the copying accuracy at longer genetic distances (Appendix A4, (Jewett *et al.*, 2012)). For samples that are sufficiently old, there is a diminishing benefit for generating larger modern reference panels (Appendix A4), which primarily results in improvements in imputation and phasing for modern samples due to recent relatedness (Jewett *et al.*, 2012; McCarthy *et al.*, 2016).

Our exploration of the impact of population demography (particularly population growth) and our empirical analysis of the male X chromosome paints a more optimistic picture for the analysis of human ancient DNA using the haplotype-copying model. We find that there is a substantial attenuation of the increase in the haplotype-copying jump-rate 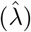 under scenarios of recent growth, and even potential decreases in the case of instant population growth (Figure 6). Together with our empirical result of the jump rate decreasing as a function of time across male X chromosomes in ancient European samples (Figure 7), the results support the idea that we may be able to impute common variants relatively accurately in human populations that have undergone recent rapid growth. Indeed the empirical accuracy of imputation is relatively high for samples within the past ~6000 years (Gamba *et al.*, 2014; Martiniano *et al.*, 2017)

As caveats, our theoretical results here do not account for some important features of aDNA data. Specifically, we have not attempted to model genotyping error and low-coverage data, both common in the analysis of ancient DNA (e.g. Dabney *et al.*, 2013). Our results on pair-wise loci could be extended to directly model the effects of errors at one or both loci. Methods using haplotype-copying HMMs with emission probabilities directly modeling low-coverage sequencing data (e.g. Rubinacci *et al.*, 2020) are more applicable to account for this sparsity in ancient DNA analysis. Another caveat is that due to the wide temporal range and the absolute number of samples available (Olalde and Posth, 2020), our empirical analyses focused on samples from western Eurasia. As ancient DNA technology improves and sampling becomes less centered on western Eurasia, it will be interesting to re-analyze the relationship between the jump-rate and sample age across multiple regions with varied demographic histories.

With the abundance of ancient DNA data being generated across a wide array of organisms, statistical and theoretical advances will need to similarly account for this new dimension in the data. Here we have highlighted the impact of time-stratified sampling for two related models, the two-locus coalescent with recombination and the haplotype copying model. We expect that our theoretical treatment of these models will serve to inform advances in statistical population genetic methods that account for serially sampled data to maximize their utility for inference.

## 4 Methods

### 4.1 Coalescent simulations and calculation of pairwise-differences

We used msprime (Kelleher *et al.*, 2016) to perform all coalescent simulations used throughout the paper. For simulations of two loci, we used a customized recombination map to reflect two non-recombining loci of a given size separated by a specified absolute recombination rate. For the simulations of haplotypes, we use the default simulation method and a uniform recombination map (default *r* = 10^−8^ per-basepair per-generation). To calculate a pairwise-coalescent effective *N_e_* to compare our constant-population-size theory for two loci with simulations under varying demographic history, we took a Monte-Carlo approach using 10^4^ coalescent simulations to compute the mean marginal pairwise coalescent time 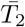 from simulations and compute 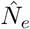 as 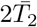.

#### 4.1.1 Monte-Carlo Simulation of Correlation in Pairwise Differences

To verify our comparisons of the theoretical prediction of *Corr*(*π_A_, π_B_*) with data, we simulated two loci as described above with a mutation rate *θ* = 0.4 (approximately equivalent to a one kilobase window with human scale parameters) for 100 log-spaced points from *ρ* ∈ [10^−4^, 10^2^]. When estimating *Corr*(*π_A_, π_B_*), we conducted 100000 independent simulations and estimated the Pearson correlation using the `pearsonr` function in the `scipy` package (Virtanen *et al.*, 2020). The standard error of the correlation was calculated using the asymptotic formula: 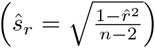.

For estimating the correlation in pairwise differences, we simulated 20 replicates of 20 Mb haplotypes and calculated a Monte-Carlo estimator of the mean correlation in segregating sites at different recombination distances. The estimation proceeds as follows: (1) we split the chromosome into non-overlapping windows of length *L* basepairs (default: 1 kilobase); (2) for each of 5000 Monte-Carlo samples we choose a window *S_A_* and define a paired window a recombination distance *r* from it (randomly choosing the direction to search); (3) compute the empirical Pearson correlation coefficient of the number of pairwise differences *Corr*(*π_A_, π_B_*) across the 5000 paired windows. Standard errors were computed using the asymptotic formula above, using the 20 replicate chromosomes. For estimation with the real whole-genome sequencing data, we use 30 logspaced bins over the range *r* = (10^−5^, 10^−3^), where *r* is in Morgans to calculate Monte-Carlo estimates of the correlation in pairwise differences. Unless otherwise specified in the text, error bars reflect two standard errors from the mean. When translating from years to generations for comparison of models to our theoretical predictions, we use a generation time of 30 years per generation from Fenner (2005).

#### 4.1.2 Monte-Carlo Estimation of Joint LD

To estimate the product of LD across timepoints (Equation 5), we used Monte-Carlo simulations of 500 modern and ancient haplotypes in a model of constant population size of *N_e_* = 10^4^. We conducted 10 replicate simulations of 1 megabase haplotypes with the mutation rate and recombination rate set to 10^−8^ per basepair per generation. We applied a filter of the minor allele frequency pooled across timepoints at *>* 5% when calculating the joint LD coefficient. We additionally bin by genetic distance using the automatic histogram binning in scipy (Virtanen *et al.*, 2020). For very low values of *ρ*, there are too few mutations co-occuring at such short distances in our simulations so we set a lower-bound of *ρ* = 1 when plotting Figure 5.

### 4.2 Analysis of Ancient Whole-Genome Sequencing Data

For our analysis of whole-genome ancient DNA data, we compared single nucleotide variants observed in the *LBK* and *Ust-Ishim* samples (Lazaridis *et al.*, 2014; Fu *et al.*, 2014). Variants were called using `samtools mpileup -C50` and were subsequently filtered using the same criterion as in DE Barros Damgaard *et al.* (2018).

To account for not having resolved haplotypes in the ancient samples, we scale the observed differences by the probability that they would be observed in a haplotype randomly sampled from the diploid genome (e.g. 0.5 if heterozygote in ancient sample, 1 if opposing homozygote in the ancient sample). For modern samples, we used haplotypes from the 1000 Genomes Project Phase 3 Dataset (Auton *et al.*, 2015).

We computed the correlation in pairwise differences in non-overlapping one kilobase windows and applied a mappability mask to account for varying coverage in the modern sample by normalizing (Auton *et al.*, 2015). Standard errors were estimated using a non-parametric bootstrap across 22 autosomes. To compare two empirical curves of 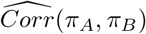, we apply a two-sided Binomial sign test to test the proportion of recombination distance bins for which one ancient sample has a higher correlation and test against the null hypothesis that the proportion is 0.5.

### 4.3 Parameter estimation in the Haplotype-Copying Model

We implemented a version of the Haplotype-Copying Model proposed by Lawson *et al.* (2012) that accounts for the genetic map distances between subsequent single-nucleotide polymorphisms. The Hidden Markov Model (HMM) is defined as follows. The transition probabilities between hidden states, *X_l_*, where *X_l_* represents the haplotype in the panel that the test haplotype copies off of at site *l*:

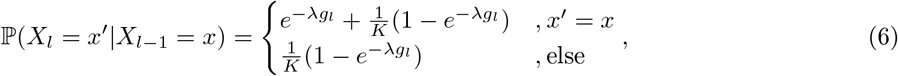

where *g_l_* is the *genetic distance* between markers *l* − 1 and *l* (in Morgan), *K* is the size of the haplotype reference panel, and *λ* is the “jump rate” or rate at which the model transitions between the haplotype copying states.

The emission probabilities can be similarly characterized, using a parameter *E* that represents the probability of a copying error:

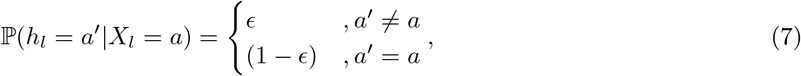

where *h_l_* is the allelic state of the query haplotype at site *l*.

We use two-dimensional numerical optimization from `scipy.optimize` (Virtanen *et al.*, 2020) to jointly estimate the maximum-likelihood estimates 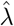 and 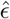. Unless specifically stated, we use the joint parameter estimates in our results for both simulated and empirical data. For profile maximum-likelihood estimates of 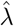, we use Brent optimization within the range [0, · · ·, 10^6^] with a fixed *E* = 10^−2^. We estimate standard errors for 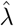 and 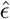 using a finite-difference approximation to the second derivative of the joint log-likelihood surface.

All simulations under the haplotype copying model were conducted using chromosomes of 40 megabases, and recombination and mutation rates of 10^−8^ per basepair per generation. Every modern panel consisted of *K* = 100 haplotypes (unless otherwise specified). We also ascertained to variants with a minor allele frequency *>* 5% in the modern panel.

### 4.4 Analysis of male X-chromosomes in 1240K Human Ancient DNA Dataset

The human ancient DNA data that we used for our analysis of the haplotype copying model (see Online Resources) are typed at a set of 1,233,013 sites across the genome and downloaded from the David Reich Laboratory’s website. Genotypes are drawn using psuedo-haploid sampling based on the available reads at these sites. We filtered the data based on the following criteria for our analysis while restricting to the X chromosome: 1) Must be a male sample; 2) Samples must not have a significant amount of modern DNA contamination (e.g. “PASS” contamination checks); 3) Samples must have ≥ 8000 non-missing variants across the X chromosome. Following this filter, the median autosomal coverage for the remaining samples is 2.303×, and an average of 1.29 sites per 25 kilobases on the X-chromosome.

Following these filters, we have a total set of 798 samples for which we estimated the maximum-likelihood jump rate under the haplotype copying model. To minimize confounding via spatial variables, we chose a centroid location (48 degrees N latitude, 6 E longitude) and only retained samples within 1500 kilometers of this centroid. Following this filtering step, there are 344 samples that are used for the main figures (Figure 7). We performed estimation of the haplotype copying jump rate across all of the 798 originally filtered samples using three different haplotype reference panels (49 CEU haplotypes [“CEU”]; 240 EUR haplotypes [“EUR”]; 1233 haplotypes [“FULLKG”]) for the X-chromsome from the 1000 Genomes Phase 3 dataset (Auton *et al.*, 2015). In all cases we used the sex-averaged recombination map for the X-chromosome from Kong *et al.* (2010). For linear modeling of the jump-rate as a function of the sample age we used the *OLS* function of `statsmodels` package (Seabold and Perktold, 2010). When comparing the real data against simulations under the demographic models inferred by Tennessen *et al.* (2012) and Browning and Browning (2015), we use *n* = 49 modern day CEU haplotypes and sampled haplotypes at ages corresponding to the real data using a generation time of 30 years per generation (Fenner, 2005). We additionally scaled each demographic model by 3*/*4 to reflect the reduced effective size of the X-chromosome.

## 5 Acknowledgements

We would like to thank all members of the Novembre, Steinücken, and Berg Labs for thoughtful feedback on this work. Particular thanks to Maryn Carlson, Harald Ringbauer, Joe Marcus for detailed discussions on earlier versions of this manuscript and Yilei Huang, and Dr. Amy Williams for detailed discussions on the results of the haplotype copying model. We thank Dr. Sharon Browning for sharing the estimated demography for the UK10K samples from their paper. We additionally thank the original study authors for sharing their data publicly, and the David Reich Lab for compiling and making publicly accessible a compilation of those data via the Allen Ancient DNA Resource (see Table S1 for detailed citations for each of the 344 ancient European samples that were used). Arjun Biddanda was supported by NIH T32 GM07197 and by NIH grant RO1HG007089 to John Novembre. This work was completed using resources provided by the University of Chicago’s Research Computing Center. The authors affirm that all data necessary for confirming the conclusions of the article are present within the article and available in a public repository (see Online Resources).

## 6 Online Resources

- Figure & Analysis Repository: https://github.com/aabiddanda/aDNA_LD_public
- Publicly-available human aDNA data from the Allen Ancient DNA Resource, compiled by the David Reich lab (v42.4 - accessed June 3rd, 2020): https://reich.hms.harvard.edu/allen-ancient-dna-resource-aadr-downloadable-genotypes-present-day-and-ancient-dna-data
- 1000 Genomes Phase 3 X Chromosome Data:http://ftp.1000genomes.ebi.ac.uk/vol1/ftp/release/20130502/
- Publicly available recombination maps: https://www.well.ox.ac.uk/~anjali/AAmap/maps_b37.tar.gz

## Appendices

## A1 The Two-Locus Ancestral Process with Population Continuity and Ancient Sampling

We first begin with a model of constant population size and where we sample one haplotype from the present and one haplotype at time *t_a_* ago (in coalescent units). The population is assumed to be constant in size with population scaled recombination rate *ρ* = 4*N_e_r*. Since we have two-samples from different time-points, we have two phases of the process: (1) where only the modern lineage can evolve at two loci (*t, t_a_*) and when both haplotypes are available to coalesce and recombine with one another (*t ≥ t_a_*). The states and possible transitions (with their corresponding rates) are shown in Figure A1.

**Figure A1:**
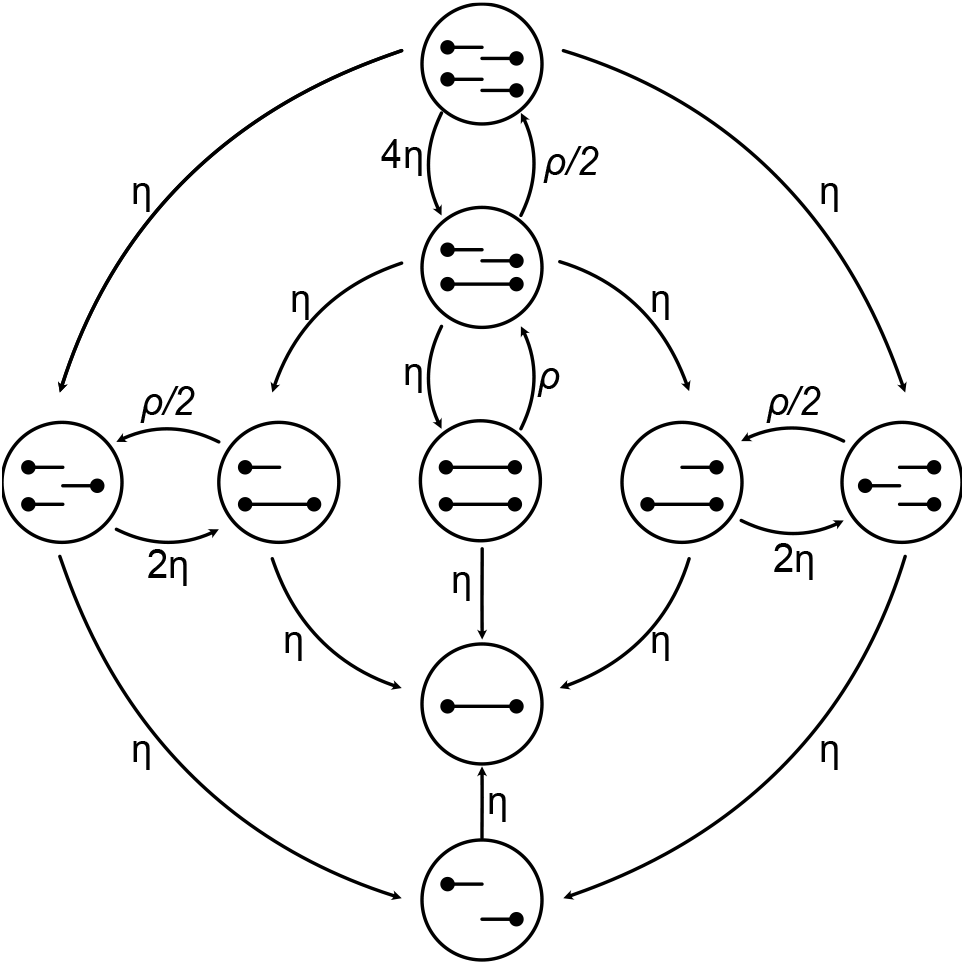
Markov chain model for the ancestral process at two loci from Simonsen and Churchill (1997). In all settings for two modern haplotypes we assume that we start from the state in the middle (state “0”) in all applications, which means that all sampled haplotypes are coupled. The parameter *η* represents the coalescent rate and the parameter *ρ* represents the recombination rate (measured in coalescent units). Figure adapted from Hobolth and Jensen (2014).

**Figure A2:**
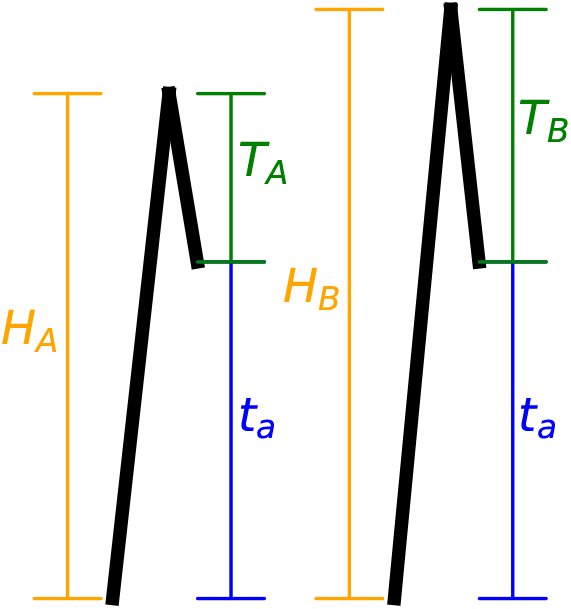
Description of variables in the two-locus case. *H* is the total tree height, *T* is the coalescent time of the ancient and modern lineage, and *t_a_* is the sampling time of the ancient lineage (in coalescent units). Here subscripts *A, B* denote the two loci separated by scaled recombination distance *ρ*.

Before calculating *joint* moments of the genealogical properties across two loci, we calculate marginal moments at individual loci: (1) *E*[*T*], the time to coalesce between the two sequences after both are able to coalesce, (2) 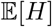, the height of the genealogy at a single locus, and (3) 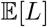, the expected total branch length at a single locus. All of these quantities are scaled by twice the population size (2*N_e_*), which we refer to as the “coalescent scale” (see Figure A1 for a schematic of these marginal quantities). The variable *T* Exponential(1) when both haplotypes are sampled from the same population. These marginal quantities can then be obtained in the model with time-stratified sampling as:

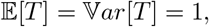

for the expectation and variance of *T*,

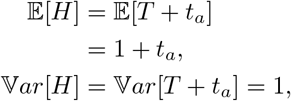

for the expectation and variance of *H*, and

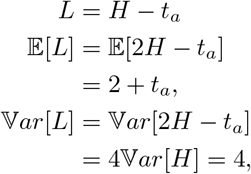

for the expectation and variance of *L*. Following the definition of these marginal moments, we calculate the covariance in the branch lengths at each locus, ℂ*ov*(*L_A_, L_B_*), as:

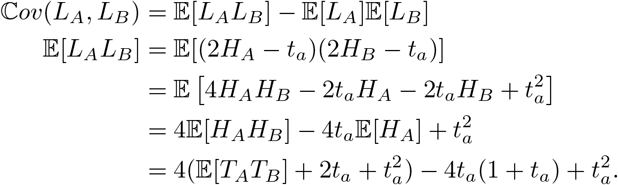

These derivations show that we can compute ℂ*ov*(*L_A_, L_B_*) under the time-staggered sampling model by computing 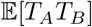.

We approach this using a “staggered” version of the Simonsen-Churchill Model as described in the main text (Simonsen and Churchill, 1997; Hobolth and Jensen, 2014) (Figure A1). In the phase where *t < t_a_*, with a single modern haplotype, we consider this as a two-state continuous-time Markov process with the rate matrix:

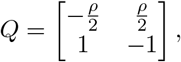

which we use to solve for the probability that the ancestral process is in state *x* at time *t_a_* as:

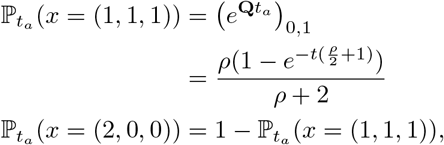

where the state *x* = (2, 0, 0) represents 2 lineages that are ancestral to both locus *A* and locus *B* and the state *x* = (1, 1, 1) represents 1 lineage ancestral to both locus *A* and *B*, one lineage ancestral to locus *A*, and one lineage ancestral to locus *B* (Hobolth and Jensen, 2014; Simonsen and Churchill, 1997). This corresponds to our “uncoupled state” in the main text. The two states in the Markov process with a single present haplotype can only be “coupled” ((2, 0, 0)) or “uncoupled” ((1, 1, 1)).

Returning to our computation of 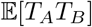 in the second phase of the ancestral process (*t > t_a_*), we obtain:

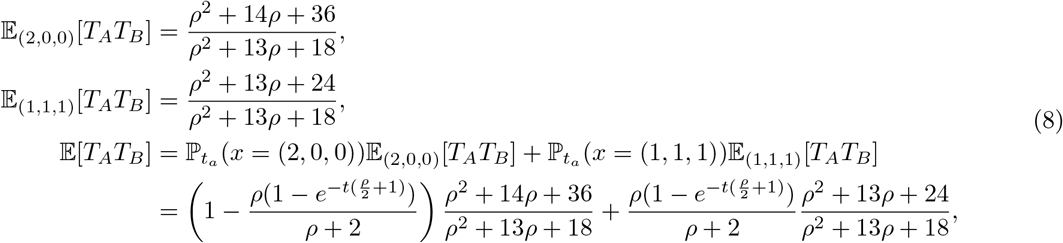

where 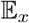 indicates the expectation conditional on starting in state *x* of the ancestral process. The first two expressions above are derived in Durrett, 2008, Chapter 3, where both haplotypes are sampled at present. The last expression is a weighting of the expectations from different starting states in the two-locus ancestral process, where the weight corresponds to the probabilities that the modern haplotype is uncoupled at the time the ancient haplotype is sampled, *t_a_*. From this we can compute the covariance in the branch length, ℂ*ov*(*L_A_, L_B_*) and ℂ*orr*(*L_A_, L_B_*): by substituting the Equation 8 into the relevant expressions previously defined, leading to the expression:

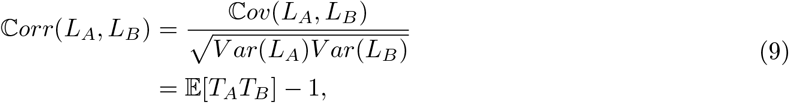

which simplifies to Equation 2 in the main text. The lower and upper limits of *t_a_* are 0 and ∞, and we show the asymptotic behavior of *Corr*(*L_A_, L_B_*) in terms of *ρ*:

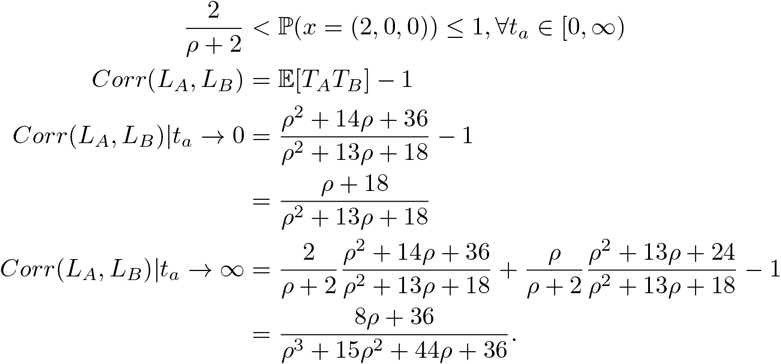

This derivation highlights the change in the rate of decay in the correlation of the branch length as a function of the sampling time from 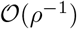 to 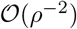.

To relate the correlation in total branch length to the correlation in the number of pairwise differences between two sequences, we use the following identities for the case where mutations occur as a Poisson process with rate *θ/*2 along branches, where *θ* is the population-scaled mutation rate (*θ* = 4*N_e_μ*) (Hobolth *et al.*, 2019):

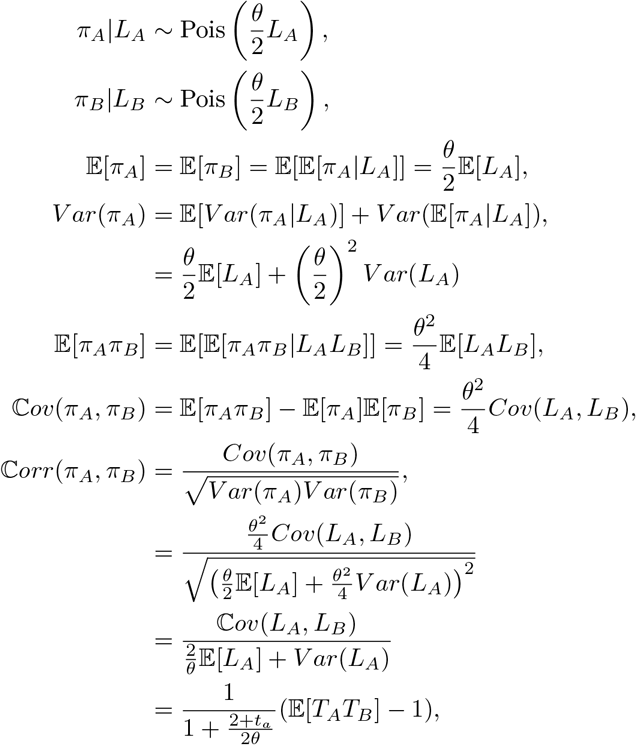

leading to a relationship with the correlation in the branch length at each locus, ℂ*orr*(*L_A_, L_B_*):

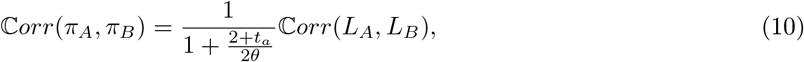

which is Equation 2 in the main text.

## A1.1 The Two-Locus Ancestral Process with population divergence and time-stratified sampling

In this section, we assume a model with divergence between the populations containing the ancient lineage and the modern lineage at the coalescent scaled time, *t_div_*. Similar to Section A1, we can partition the ancestral process into three phases: (1) when the modern lineage is the only one evolving, (2) when the ancient lineage and the modern lineage are both evolving *but are not able* to coalescent with one another and (3) when both lineages are in the ancestral population and can coalesce with each other. These three phases can be seen Figure A3.

**Figure A3:**
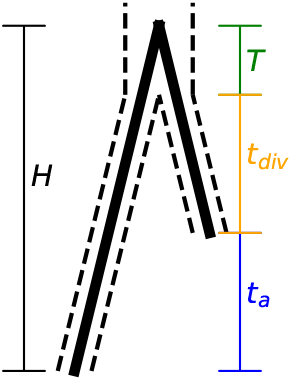
Description of variables in the single-locus case. *H* is the total tree height, *T* is the coalescent time of the ancient and modern lineage, and *t_a_* is the sampling time of the ancient lineage (in coalescent units)

The model with population divergence has an additional parameter, *t_div_*, the divergence time of the two populations. We first show the properties of the marginal tree under the divergence model (see Section A1, and Section A3, for a definition of the quantities):

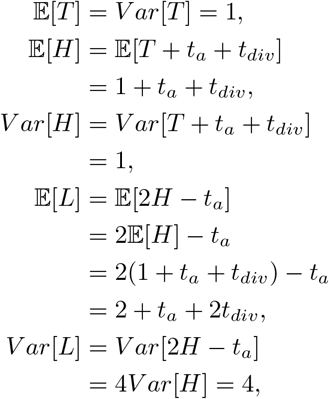

where *t_div_* is the population divergence times in coalescent units, *t_a_* is the sampling time of the ancient lineage, *T* is the exponentially distributed time after both lineages are able to coalesce that they coalesce with one another. Using these results, we can calculate moments of the joint distribution of genealogical properties like the tree height (*H*), and total branch length (*L*). Specifically, the two-locus ancestral process behaves independently within each population for time *t_a_* and *t_div_* and each population is assumed to have the same population size. We begin by deriving the joint expectation of tree-height *H_A_H_B_*:

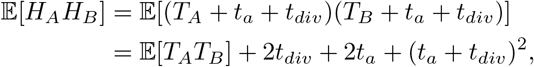

and joint tree length *L_A_L_B_*:

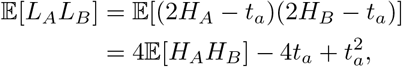

where we must solve for the joint expectation of 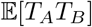, but with the additional complication of population divergence. In order to do this we must calculate the probability of being in one of three starting states at time *t_a_* + *t_div_*: (1) the state *x* = (2, 0, 0) where both the ancient and modern haplotypes are “coupled”, (2) the state *x* = (0, 2, 2) where both the ancient and modern haplotype are “uncoupled”, which is possible due to the independent evolution of both lineages during *t_a_ < t < t_a_* + *t_div_*, and (3) state *x* = (1, 1, 1) where one haplotype is uncoupled while the other is coupled. We consider the two independent processes within each population until the divergence time and calculate the probabilities of being in each starting state as follows:

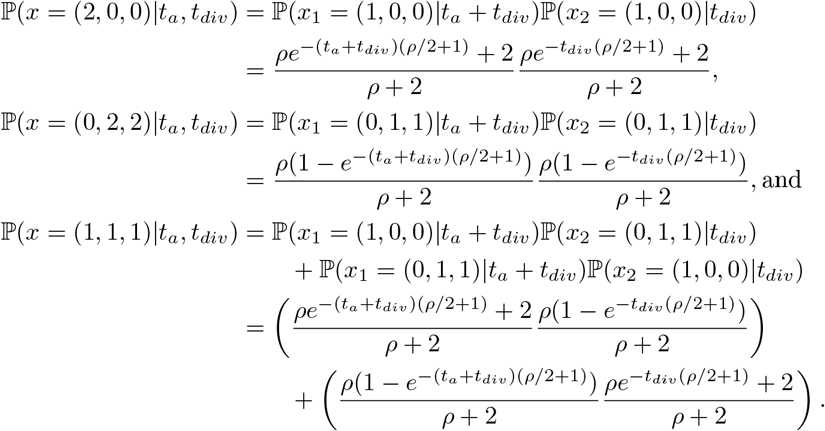

From these probabilities we calculate the expectation of the joint coalescent times conditional on being in a specified state at time *t_a_* + *t_div_* is obtained as:

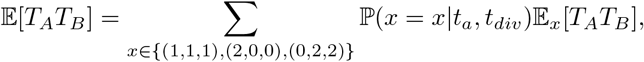

where each of 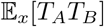 are defined using previously derived results under the two-locus ancestral process conditional on being in a starting state *x* (Simonsen and Churchill, 1997; Durrett, 2008, Chapter 3). This is different from the model under population continuity (where the *x* = (0, 2, 2) state was not possible). If we set *t_div_* = 0, then this corresponds exactly to the model without population divergence. While the underlying mathematical results are more involved, they provide insights on how population divergence affects joint coalescent times.

We can now compute joint statistics (e.g. correlation) of the tree properties at each of the loci following common formulas, for example for the correlation in total branch length at each locus:

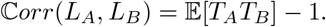

## A2 Expectations of joint coalescent times under the time-stratified model

We assume that the following results on the joint coalescent times for two contemporary haplotypes starting in the same state in the two-locus ancestral process as defined in Durrett, 2008, Chapter 3 are known:

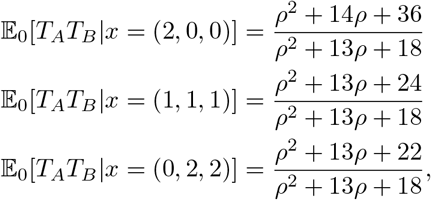

and now we will go through the individual cases for the time-stratified case: **(1)** both modern and ancient haplotypes start coupled, **(2)** both modern and ancient haplotypes are “uncoupled” and finally **(3)** where *only one* of the modern and ancient haplotypes are coupled (the other is uncoupled).

We first define two quantities, called *γ* and *η*. The variable *γ* refers to the probability of starting in the coupled ((1, 0, 0)) state and ending in the uncoupled state ((0, 1, 1)) at time *t_a_* for a single haplotype (which is Equation (1) in the main text). The variable *η* is the converse, the probability of starting in the uncoupled state and ending in the coupled state at time *t_a_*. Using the matrix exponential 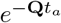 of the following rate matrix for the process with a single haplotype:

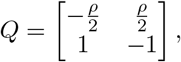

we arrive at the following expressions for *γ* and *η*:

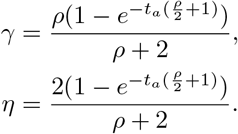

With these in hand we can start tackling our first case **(1)** from above:

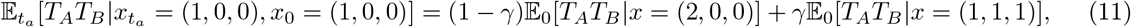

where, *x*_0_ = (1, 0, 0) indicates that the modern haplotype is coupled, and 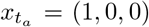 indicates that the ancient haplotype is coupled as well. This holds because the modern haplotype can be coupled with probability 1 − *γ* leading to state *x* = (2, 0, 0) for the joint ancestral process, or it can be uncoupled with probability *γ* resulting in state *x* = (1, 1, 1). For case **(2)**(both haplotypes uncoupled) we obtain:

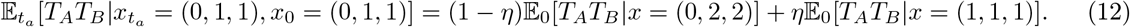

The final case **(3)** is the most complicated and we break this into a further two sub-cases below:

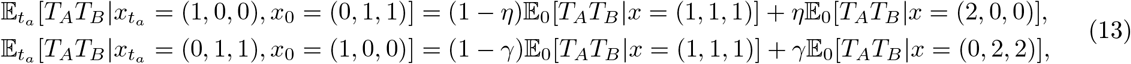

where the first case corresponds to the modern haplotype starting in the “uncoupled” state (denoted by the *x*_0_ in the expectation) and the second case corresponds to the modern haplotype starting in the “coupled” state.

## A3 The expected product of linkage disequilibrium between time-stratified samples

Here, we derive the scaled product of linkage disequilibrium between time-stratified samples normalized by the heterozygosity across both sites and time points. We first start from the definition of the statistic in terms of haplotype and allele frequencies in the ancient and modern samples:

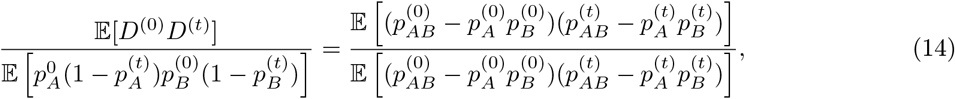

where 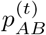 is the frequency of the haplotype with the derived alleles at both loci at time *t*, 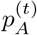 is the frequency of the derived allele at the first locus, and 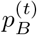 is the frequency of the derived allele at the second locus. Using the approach of McVean (2002) we define this ratio using branch lengths in the genealogy relating certain modern and ancient samples, where a mutation would result in a observed pattern of identity by state. We first expand the numerator as follows:

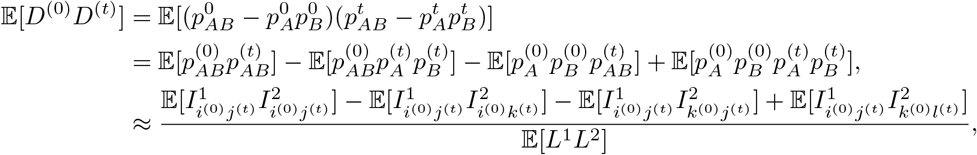

where *i, j, k, l* denote sampled haplotypes. Furthermore, 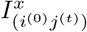 is the branch length leading from the *T_mrca_* of the samples *i*^(0)^ at time 0 and *j*^(*t*)^ at time *t* to the *T_mrca_* of the total population (including the ancient individuals) at locus *x*. 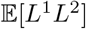 is the joint expectation of the total genealogical branch length for the complete population at both loci. The approximation in the final step above follows from assuming a small mutation rate (McVean, 2002). We use the definition 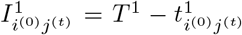, where *T* ^1^ is the *T_mrca_* for the total population (modern and ancient) at the first locus and 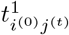 is the pairwise coalescent time for samples *i*^(0)^*, j*^(*t*)^ at the first locus. Using this relationship between coalescent times and identity coefficients, we arrive at:

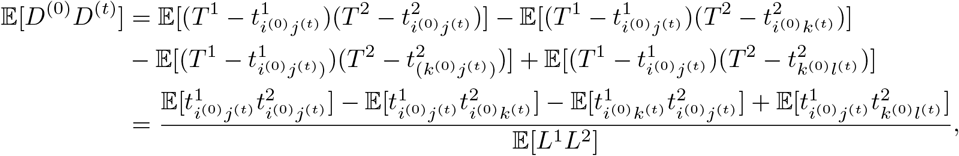

where we have used the argument that terms of the product of a pairwise coalescent time at one locus and the total *T_mrca_* at the other locus (e.g. 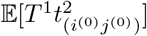 do not depend on the indices *i, j* (Durrett, 2008, Chapter 3). This means that the numerator of ^(^t^*i*^he^*j*^ex^)^pression above can be computed using the expectations of pairwise coalescent times in the time-stratified model (see Appendix A2).

The denominator of our expression 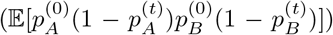 is the probability of drawing two haplotypes at the first locus that are at different time points and differ in their allelic identity, *and* drawing two haplotypes at the second locus from different timepoints that also differ in their allelic identity. This is a measure of the time-stratified joint heterozygosity at both sites. We note that this is different from the interpretation of 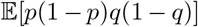 which is the probability of a difference at the first locus and a difference at the second locus under a random draw from of a sample from a contemporary population and is the denominator of 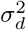 (McVean, 2002). We define the denominator similarly using pairwise coalescent times as:

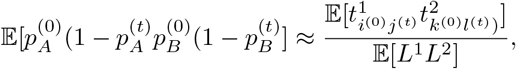

where we see that joint total branch length term 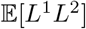 will cancel out when evaluating the ratio. We can now turn to actually computing this expression using the joint expectations for coalescent times calculated in our time-stratified model (see Appendix A2 for the derivation of these joint coalescent times):

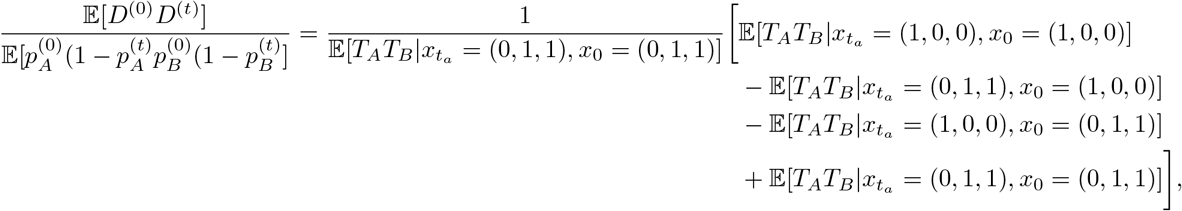

which can be simplified to the following expression after substituting the proper expressions for the joint coalescent times derived in Appendix A2:

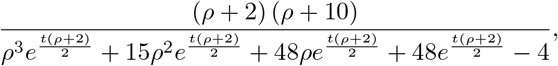

which is the expression reported in the main text (Equation 5). Importantly, we find that when *t* = 0, the expression simplifies to 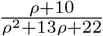 which is the expression for 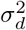 in the case with two contemporary samples (McVean, 2002).

## A4 Expected-Time to First Coalescent for an Ancient Sample

Here we consider a single ancient haplotype sampled at a time *t_a_* in the past and how it coalesces into the ancestral lineages of a reference panel of size *K* haplotypes sampled at the present. We define the random variable *T* ^∗^ as the additional time of a coalescent event involving the ancient haplotype and a lineage ancestral to the modern reference panel after the time that the ancient haplotype is sampled (*t_a_*). The expectation of this quantity can be written as:

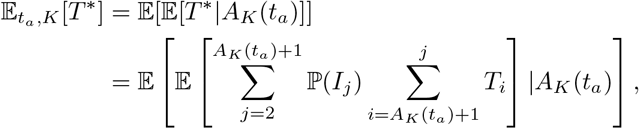

where *A_K_*(*t_a_*) is the number of lineages ancestral to the modern reference panel at time *t_a_*, ℙ(*I_j_*) is the probability that the *j^th^* coalescent event involves the ancient lineage, and *T_i_* is the *i^th^* inter-coalescent time.

Starting at time *t_a_* with *n_t_* lineages, we calculate the probability that the *j^th^* coalescent event involves the ancient lineage as:

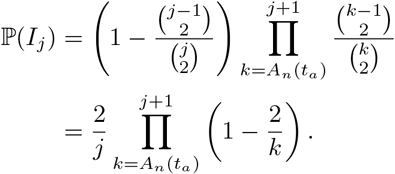

In a constant population size model, we have 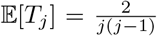. Using this fact, the expected time until the first coalescence involving the ancient lineage (*T*\*) is:

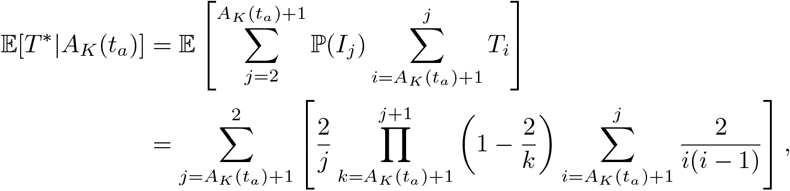

and considering the summation over *A_K_*(*t_a_*), we arrive at our final expression:

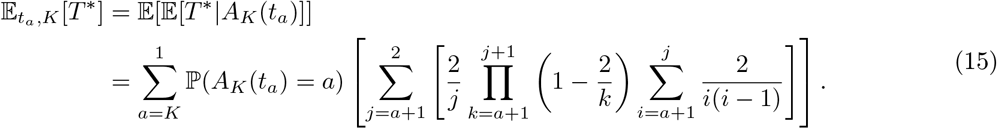

The probability distribution ℙ(*A_K_*(*t*) = *a*) involves a number of alternating sums and leads rapidly to numerical error as the sample size gets large (see Equation 15 in Chen and Chen (2013)). To alleviate this issue, following Jewett and Rosenberg (2014) we approximate ℙ(*A_K_*(*t*) = *a*) as 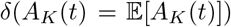. That is, rather than calculate the probability distribution of *A_K_*(*t*) across states 1*…K*, we will approximate it with its expectation 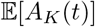. One approximation for 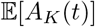 is found in Griffiths (1984):

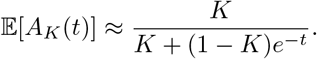

Further approximations for this expectation exist and are explored in greater detail in Jewett and Rosenberg (2014). We chose the above approximation largely for computational convenience as it does not involve any summation, has a simple form, and is comparably accurate when compared to other approximations (Jewett and Rosenberg, 2014).

The additional time to coalescence for the ancient sample 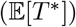 is proportional to the number of recombination events that can affect the genealogical closest haplotype to the ancient sample that is in the modern panel. For example, for a sample with *t_a_* = 2 × 10^−4^ there is 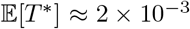 and 2 × 10^−4^ with a panel size of *K* = 1000 and 10000 respectively (Figure S4). This guides the intuition that for large panel sizes and recent sampling times, the time for the ancient haplotype to coalesce with the panel is quite small, and therefore we expect the haplotype copying rate to be fairly small (leading to longer shared blocks). This is the key intuition behind long-range phasing methods that take advantage of recent relatedness (e.g. Loh *et al.*, 2016). For samples on the order of ~ 10^−2^ coalescent units, the relative ratio is 1.17 for 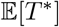 with modern panel sizes of *K* = 1000 and *K* = 10000 (as opposed to 6.99 when *t_a_* = 10^−4^). This highlights a saturation effect of within-panel coalescence at deeper times, limiting the expected utility of large modern panels for the setting with substantially ancient samples (Figure S4).

## S5 Supplementary Figures

**Figure S1:**
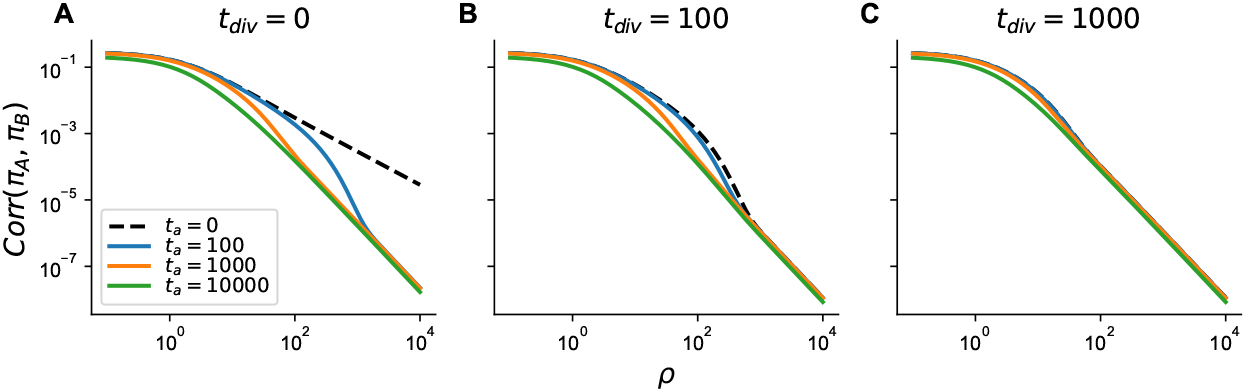
Theoretical correlation in pairwise differences for samples under a model with divergence. Divergence times *t_div_* are **(A)**0, **(B)**100, and **(C)**500 generations in the past.

**Figure S2:**
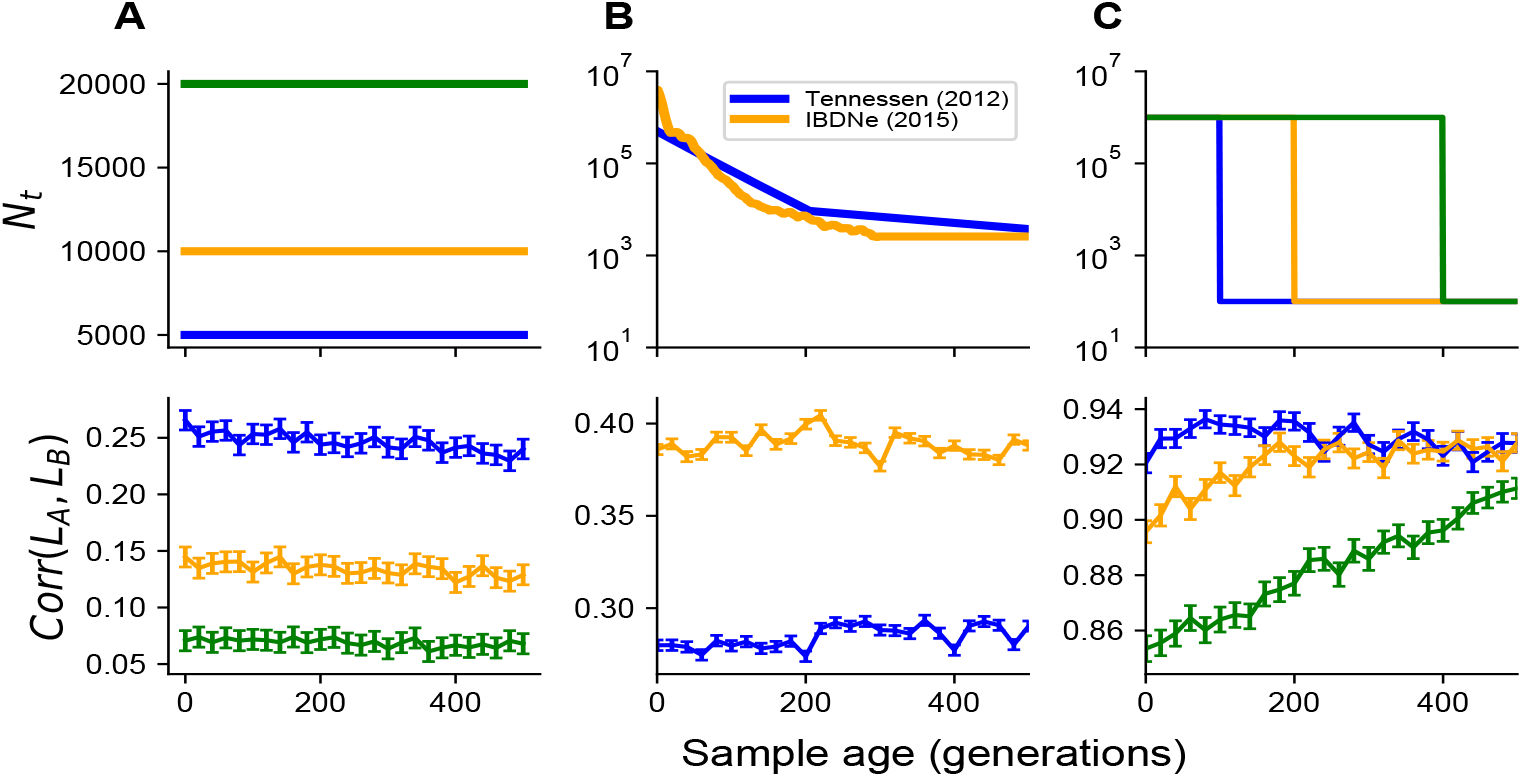
Effect of demographic history on the correlation in the total branch length between two loci. For all simulations, the recombination rate was set to 10^−4^ per generation. Simulated scenarios include: **(A)** constant population size, **(B)** inferred models of population growth, and **(C)** models of instantaneous population growth.

**Figure S3:**
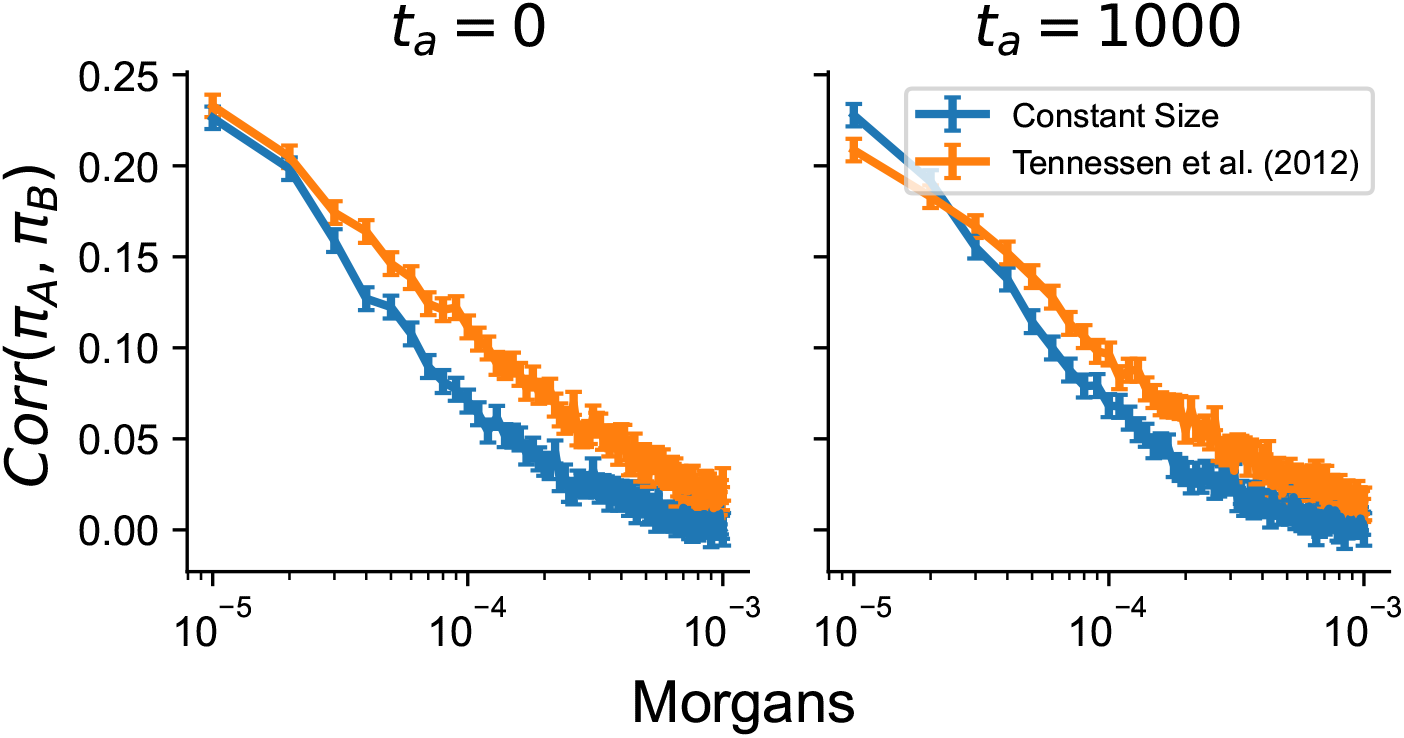
Effect of demographic history on the correlation in pairwise differences. Recombination points on the x-axis are considered as the mean recombination distance for windows of 1 kilobase separated by *d* windows (using *r* = 10^−8^ as the recombination distance).

**Figure S4:**
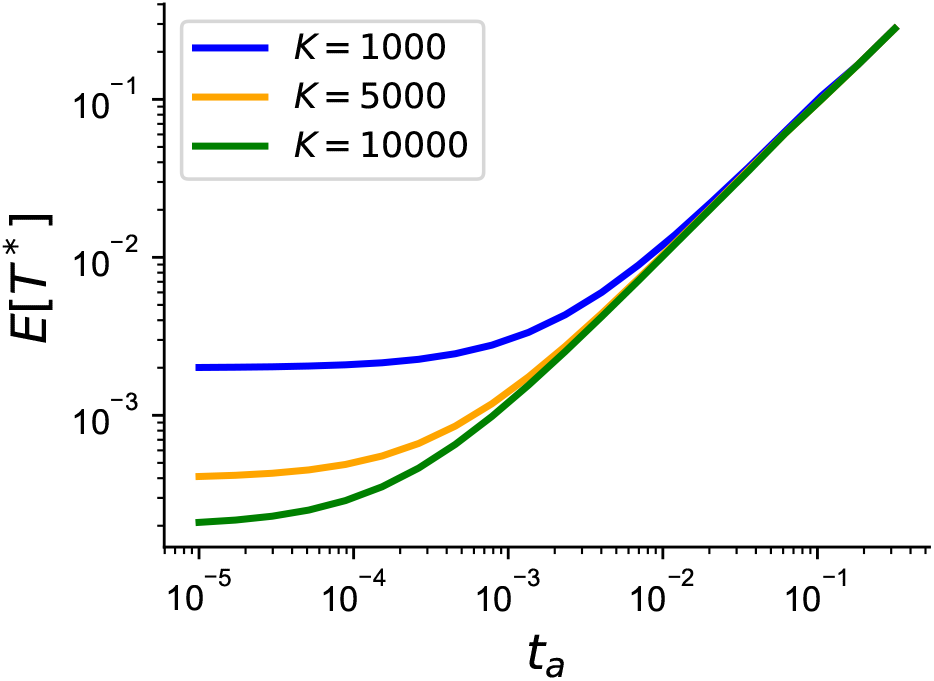
Expected time to first coalescent event involving an ancient haplotype with lineages ancestral to modern panel in a model of constant population size. As an approximation, with an *N_e_* = 10^4^, a sample with *t_a_* = 10^−3^ is 10 generations old(~300 years old (Fenner, 2005)), and there is little difference between reference panel sizes of *K* = 5000 and *K* = 10000.

**Figure S5:**
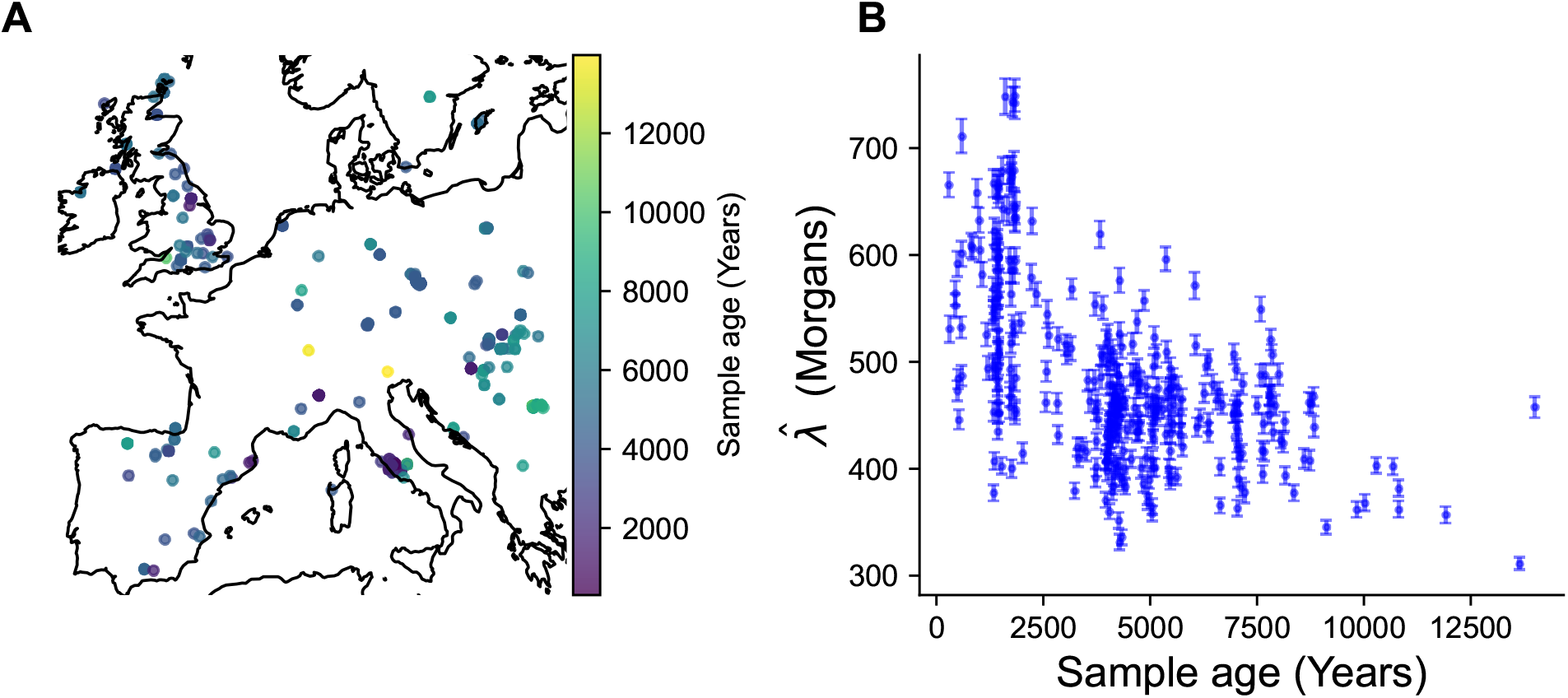
**(A)** Map of samples *<* 1500 kilometers from hypothetical location of central Europe (see Methods) **(B)** Decrease in estimated 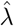 as a function of sample age in generations when estimated from male X chromosomes using all male X chromosomes from samples in the CEU population (*n* = 49) from *Auton et al.* (2015).

**Figure S6:**
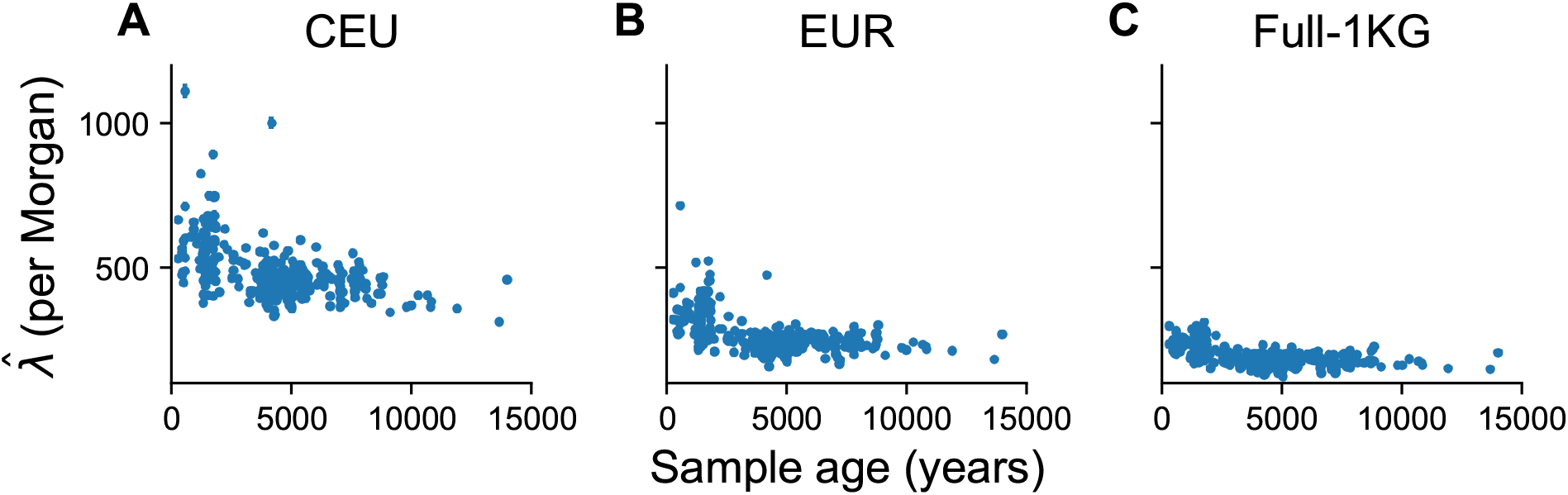
The impact of panel size on the estimated haplotype copying jump rate. **(A)** Using all male CEU individuals (n=49), **(B)** using all male individuals from the european (EUR) regional grouping (n=240), and **(C)** using all male individuals in the 1000 Genomes (n=1233). As the panel size increases, the estimated jump rate decreases in terms of the absolute scale, indicating longer shared haplotypes due to closer relatedness. However, in all cases we still find that the jump rate decrease as a function of the sample age (*p <* 0.05; linear regression). For all simulations we use the same set of samples as shown in the main text (restricting to *<* 1500 kilometers from a centroid in central europe).

**Figure S7:**
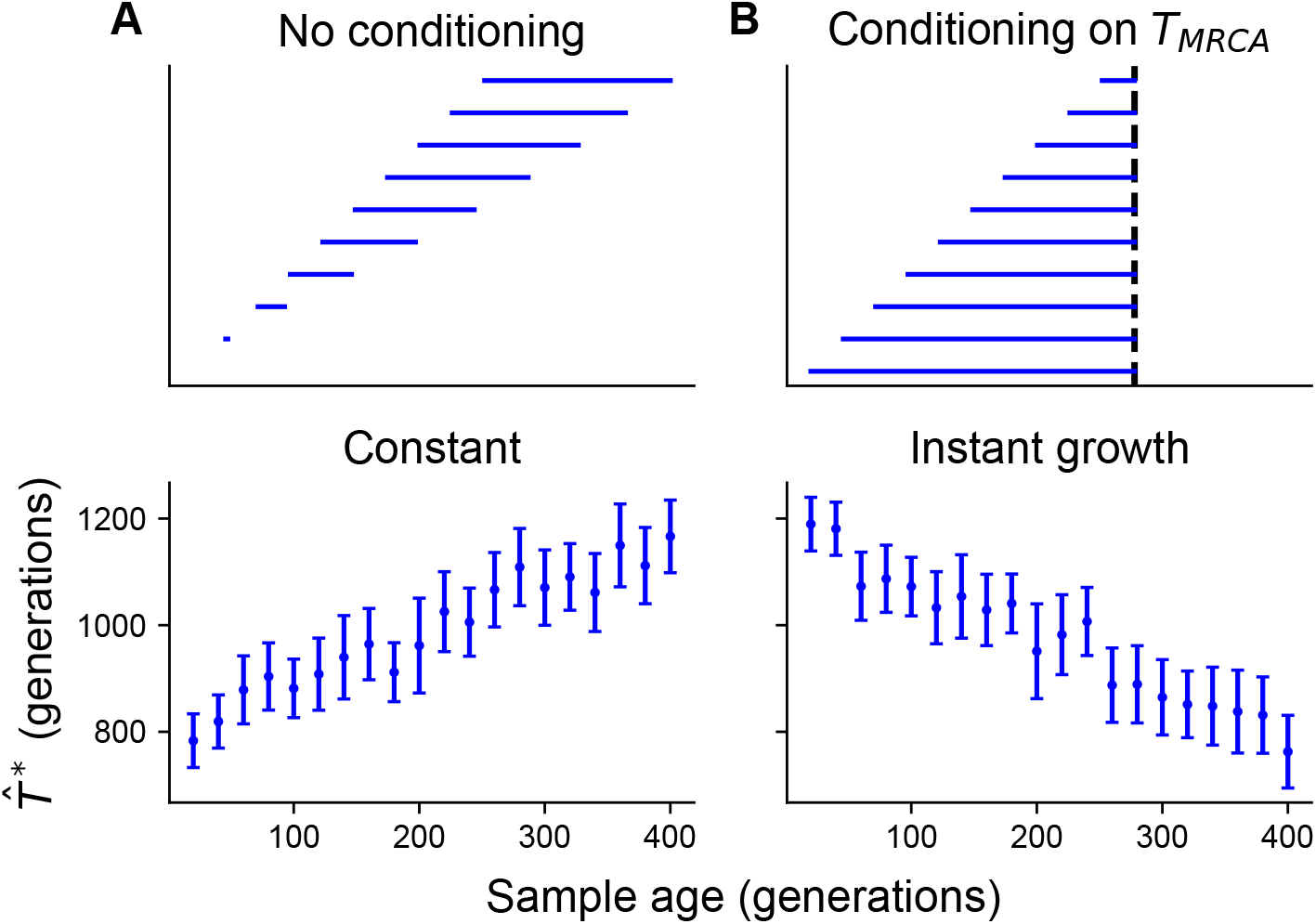
Schematic illustration of the impact of conditioning on the *T_mrca_* with simulated data. **(A)** In the case of no-conditioning, we simulate the time to first coalescence for an ancient sample with a lineage ancestral to a member of a modern panel (*K* = 100) in a constant population size scenario of *N_e_* = 10^4^. **(B)** To mimic the case of conditioning on the *T_mrca_*, we simulate instantaneous growth at 400 generations from a population size of 10^4^ to 10^6^. In both simulations, ancient haplotypes are sampled every 20 generations, are conducted using 5000 replicates and error bars represent 2 standard deviations from the mean time to first coalescence.

**Figure S8:**
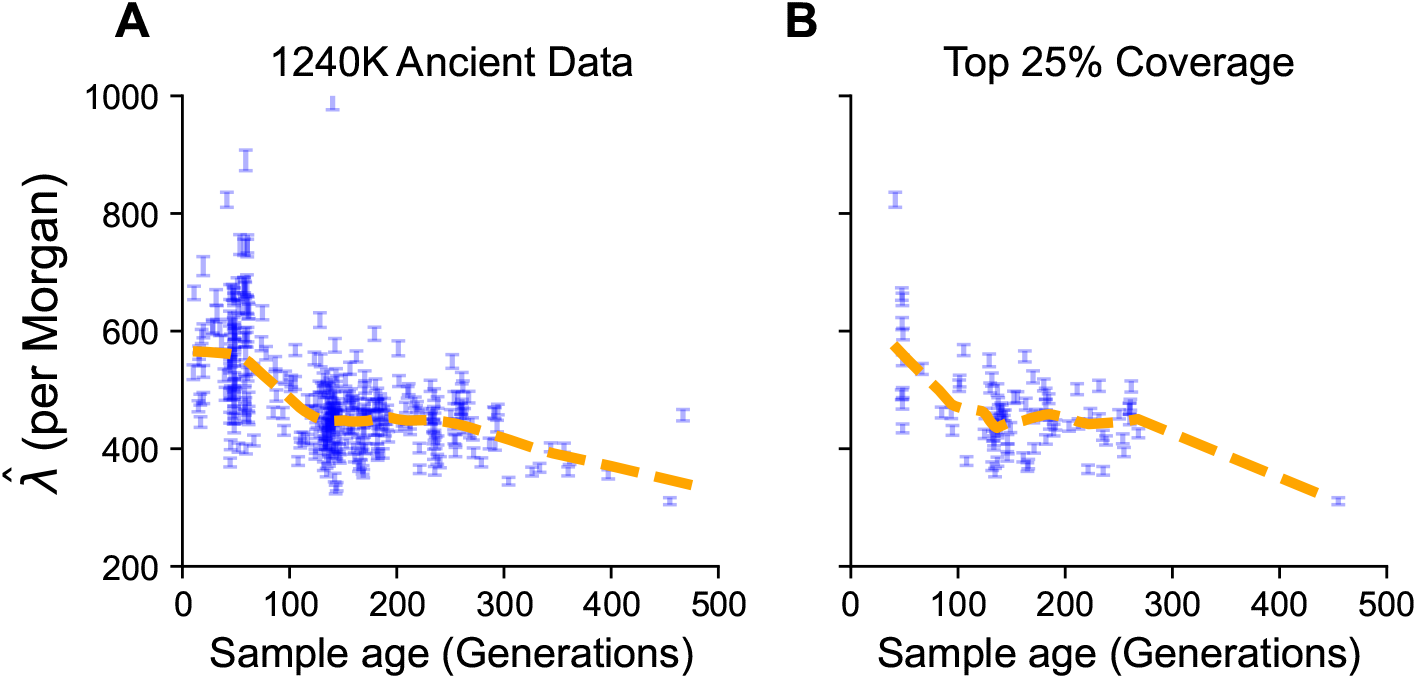
The effect of filtering by coverage on the estimated jump rate as a function of time. **(A)** The original dataset restricted to European individuals. **(A)** The evaluation of the haplotype copying jump rate restricted to individuals with the top 25% of the empirical autosomal coverage distribution. In both cases the qualitative decrease in the jump rate as a function of time is present.

**Figure S9:**
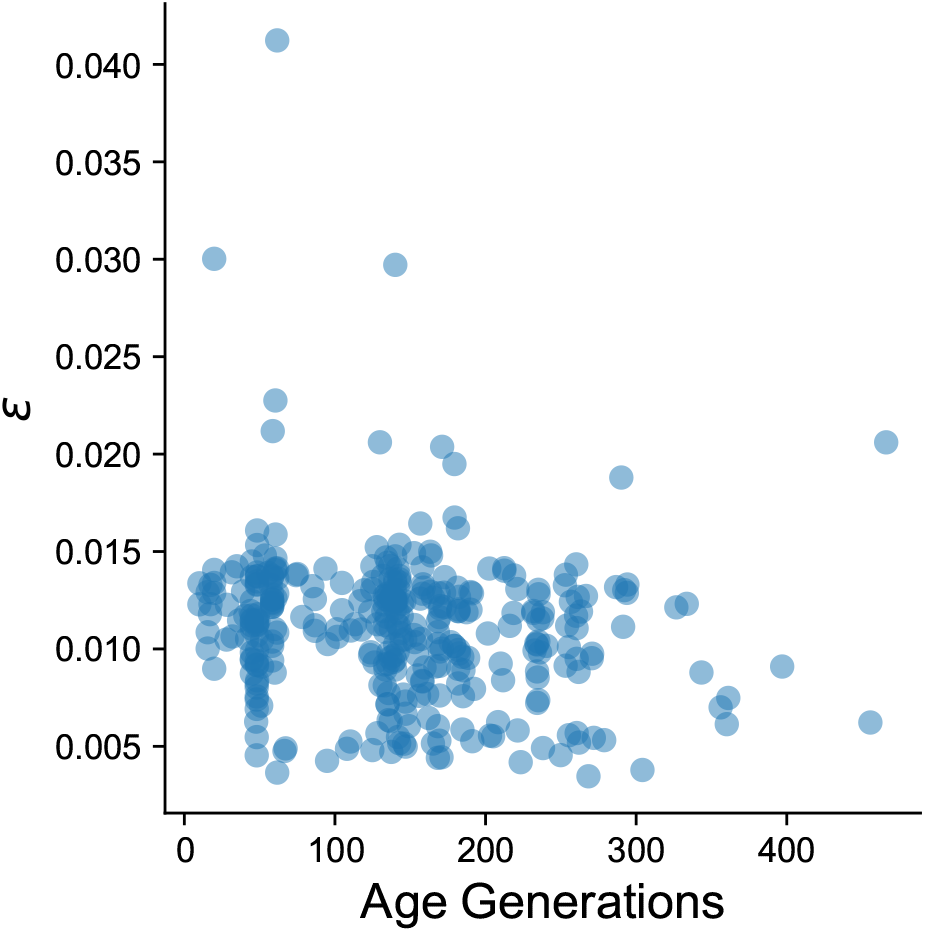
Estimated haplotype-copying error rate as a function of sample age. There is a weak anticorrelation between the estimated error rate and the sample age (*r* = −0.17, 95% CI=[−0.27, −0.06]).

## S6 Supplementary Data

**Table S1:**
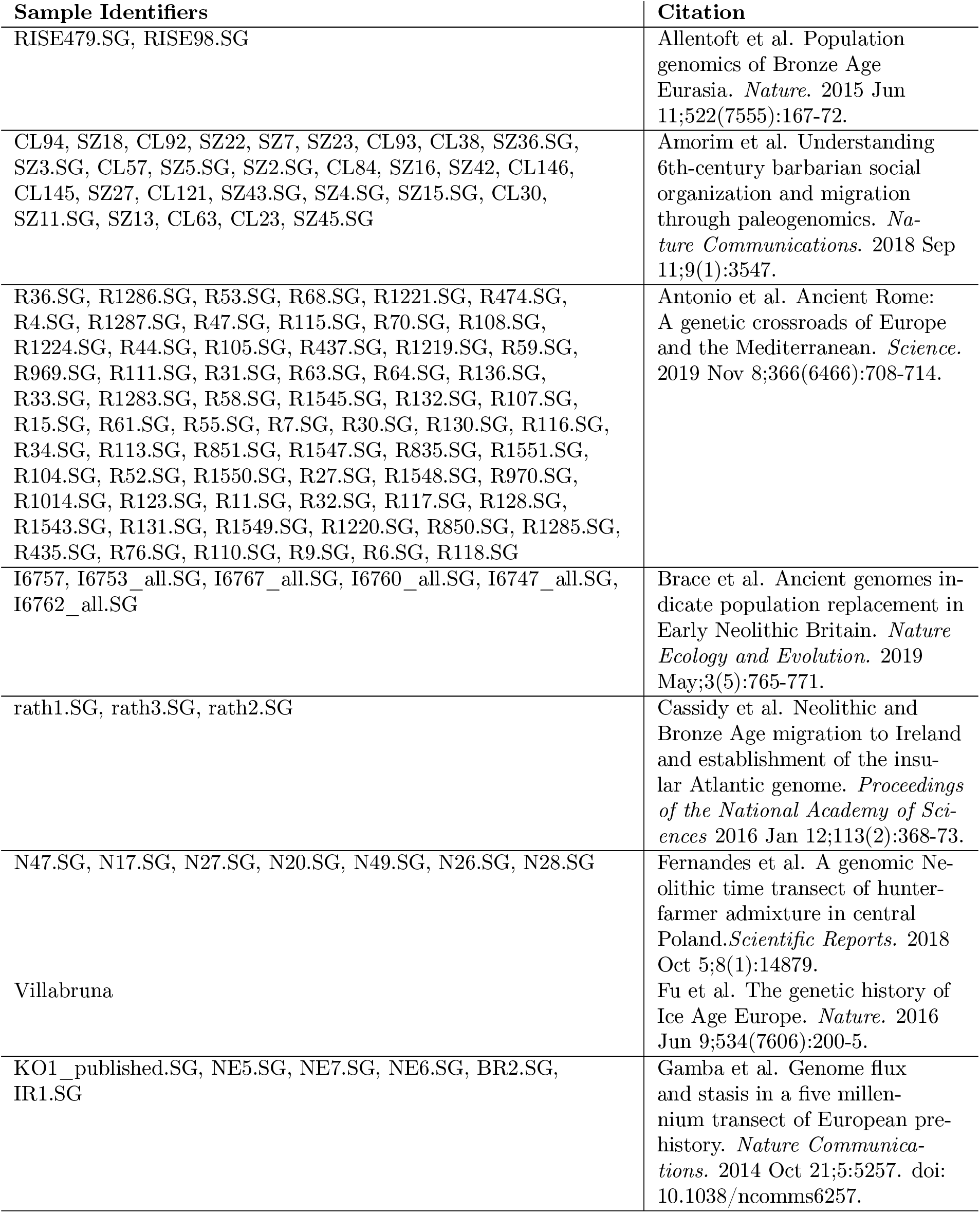
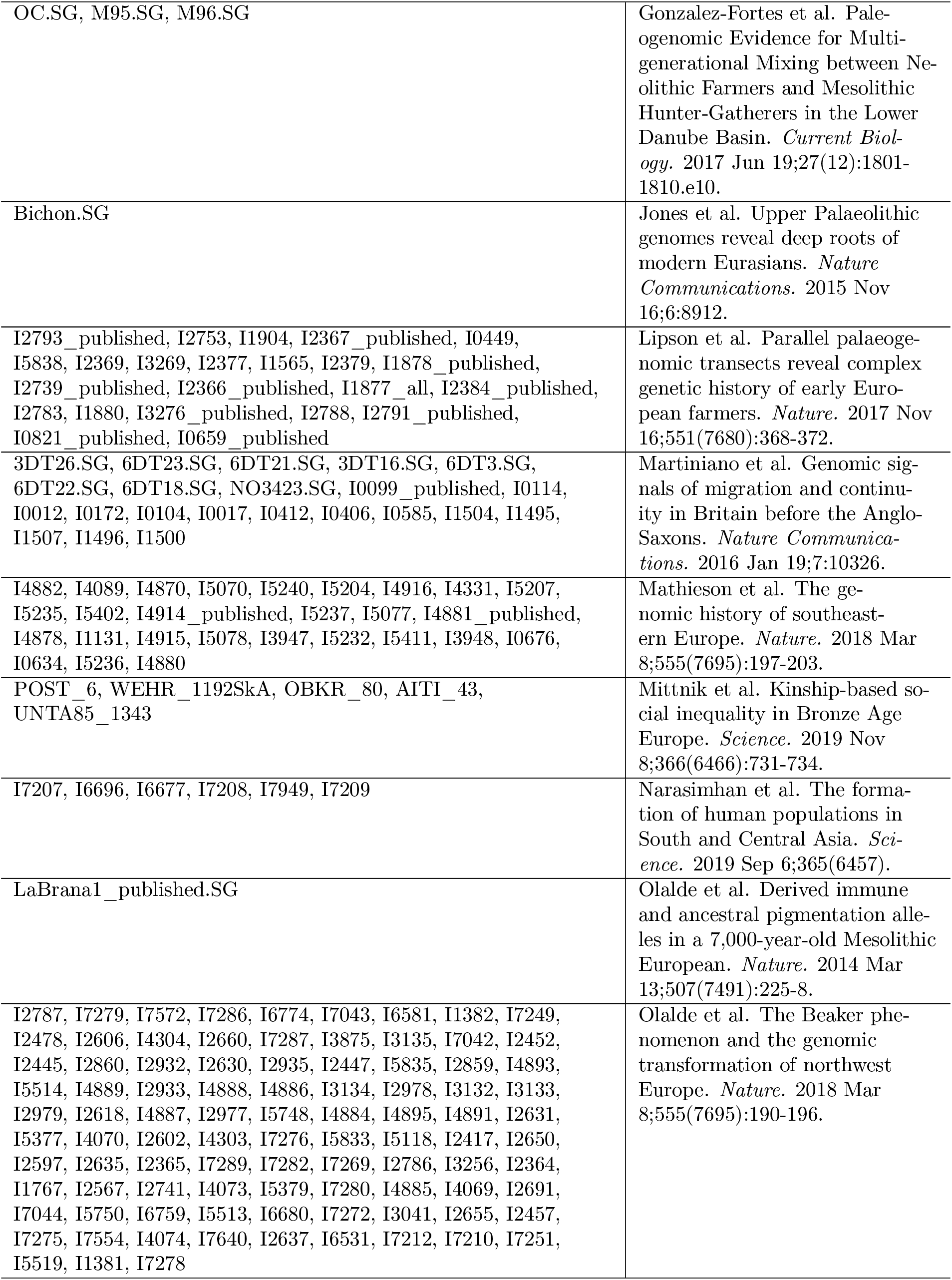
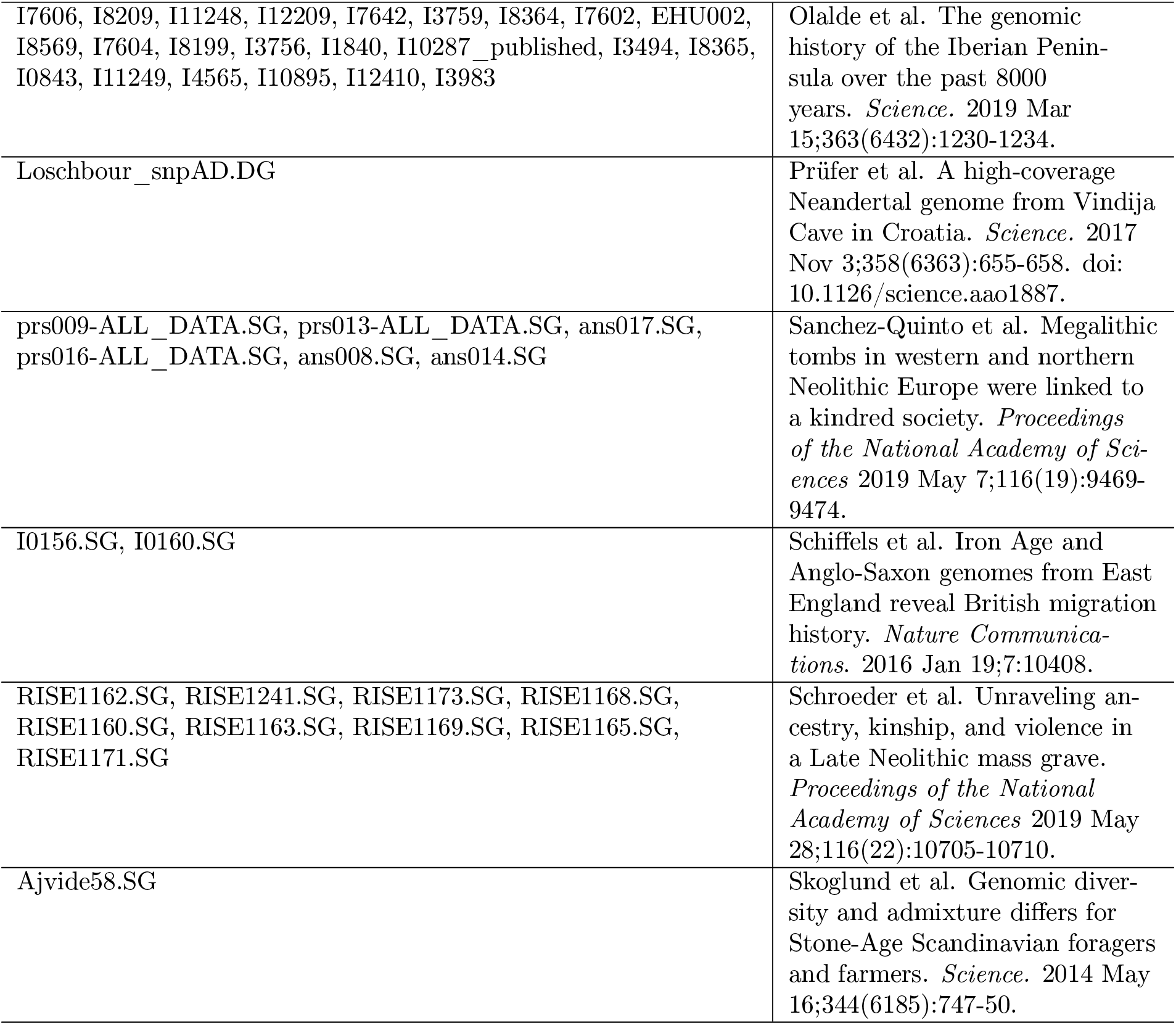
Sample identifier information and corresponding publication citation for ancient DNA samples used from Allen Ancient DNA Resource.

